# DynamicGT: a dynamic-aware geometric transformer model to predict protein binding interfaces in flexible and disordered regions

**DOI:** 10.1101/2025.03.04.641377

**Authors:** Omid Mokhtari, Sergei Grudinin, Yasaman Karami, Hamed Khakzad

## Abstract

Protein-protein interactions are fundamental to cellular processes, yet existing deep learning approaches for binding site prediction often rely on static structures, limiting their performance when disordered or flexible regions are involved. To address this, we introduce a novel dynamic-aware method for predicting protein-protein binding sites by integrating conformational dynamics into a cooperative graph neural network (Co-GNN) architecture with a geometric transformer (GT). Our approach uniquely encodes dynamic features at both the node (atom) and edge (interaction) levels, and consider both bound and unbound states to enhance model generalization. The dynamic regulation of message passing between core and surface residues optimizes the identification of critical interactions for efficient information transfer. We trained our model on an extensive overall 1-ms molecular dynamics simulations dataset across multiple benchmarks as the gold standard and further extended it by adding generated conformations by AlphaFlow. Comprehensive evaluation on diverse independent datasets containing disordered, transient, and unbound structures showed that incorporating dynamic features in cooperative architecture significantly boosts prediction accuracy when flexibility matters, and requires substantially less amount of data than leading static models.

## Introduction

Protein-protein interactions (PPIs) play a crucial role in many cellular processes and act as structural and functional components within the cell. Identifying the interaction site is crucial to understanding protein functions and serves as the foundation for protein modeling, drug targeting, and therapeutic development [1, 2]. The prediction of PPI sites has seen remarkable progress with the advent of deep learning, leveraging both sequence- and structure-based approaches. Although sequence-based methods [3–5] enable a broader applicability of the model, structure-based techniques have demonstrated superior performance. Among the latter, one important aspect of developing new models lies in determining how best to encode protein structures. A powerful framework for encoding both atom-specific features as well as the interactions between atoms is graph representations of protein structures. Graph neural networks (GNNs) has achieved state-of-the-art performance across diverse applications [6–10].

Recent advancements in GNNs, such as geometric vector perceptrons (GVP) [11] and the geometric transformer [12] have further opened novel geometric-based strategies, resulting in state-of-the-art tools such as PeSTo [13]. In addition to classical PPI prediction, recent studies have focused on the role of intrinsically disordered regions (IDRs) in mediating protein interactions [14, 15]. AlphaFold (AF) [16, 17] has revolutionized the field, but still struggles to detect correct binding interfaces when some extent of flexibility is involved such as in IDRs [18, 19] as dynamic information is absent in its training set. Utilizing a dataset of 42 protein-IDR complexes, Bret et al. [19] showed that while AF-multimer [20] only achieves 40% success rate over full length sequences, a fragmentation strategy can increase the success rate up to 89% where the fragment with highest ipTM score corresponds to the region containing the correct binding site. Interestingly, they showed that one of the difficulties encountered by AF-multimer in dealing with large IDR-containing proteins lies in its ability to identify the correct binding site within the partner proteins. While their method showed progress in this direction, for orphan proteins with no homologous sequences, such evolutionary information would be missing. In addition large protein complexes see difficulties to be fragmented to find a correct interaction site. Omidi et al. [21] also showed that AF-multimer can capture the IDR interactions while it is less successful when dealing with more heterogeneous, and fuzzy-like interactions [22].

Protein’s conformational dynamics is intricately linked to its function [23, 24]. At interaction sites, changes in solvent-accessible surface area (SASA) can significantly affect interaction specificity and binding affinity [25, 26]. Binding events often alter protein dynamics, and distant sites contribute to function through inter-residue communication pathways [27, 28]. These interconnections highlight the role of structural dynamics and the core residues in mediating protein interactions and functions. To capture the full conformational space of proteins, several approaches have emerged, including the refinement and modification of AlphaFold [29–33], alongside generative models trained on molecular dynamics (MD) data [34]. While computationally intensive, MD simulations remain as a gold standard for providing reliable, and detailed insights into protein dynamics [35].

Considering protein conformational dynamics in a deep learning framework for different downstream tasks including binding site prediction holds considerable promise, yet challenging due to several technical difficulties. For example, in graph-based representations, edge definition remains a challenge; some methods use distance cutoffs [36], while others enforce a fixed number of neighbors [13], which can be inconsistent (e.g., surface nodes should have fewer neighbors than core nodes). The choice between all-atom representations and surface-focused approaches such as MaSIF-site [37] that abstracts the core residues, remains a topic of debate, while integrating both through a so-called bipartite graph representation has proven advantageous [38], as subsequent studies demonstrated the potential importance of core residues to enhance prediction [39].

Here, we introduce the Dynamic-aware Geometric Transformer (DynamicGT), a novel GNN-based architecture designed to capture the conformational heterogeneity of proteins for residue classification in protein binding sites. Inspired from the principles of cooperative graph neural networks [40], DynamicGT enables dynamic message passing between surface and core atoms, where a geometric transformer imposes strict message-passing constraints to predict binding site probabilities. We compiled an extensive dataset of MD simulations, and expanded it with generated conformations by AlphaFlow. This dataset served as a comprehensive resource for training the model to predict protein-protein interfaces. Our findings revealed the crucial role of cooperativity between surface and core residues and the importance of dynamic features in enhancing predictive accuracy for flexible and disordered regions. By incorporating the conformational dynamics of proteins, our model outperformed existing IDR site predictors and demonstrated enhanced consistency across a wide range of bound- and unbound-state benchmarks.

## Results

### The dynamic co-operative graph transformer architecture

We developed a novel architecture that leverages protein conformational heterogeneity and incorporates differentiation of the core/surface nodes using cooperative GNNs (Figure 1a, Supplementary Figure S1-S3). This enables the model to autonomously determine and regulate the communications between surface and core residues to predict binding sites. In the first stage, a basic GNN predicts atom-level actions (listening and broadcasting). Communication constraints are then applied to surface-core interactions, while intra-type communications remain unrestricted. The masked graph is then passed to a geometric transformer, and node embeddings are pooled from atom to residue level, followed by label classification through a decoding layer.

**Figure 1:**
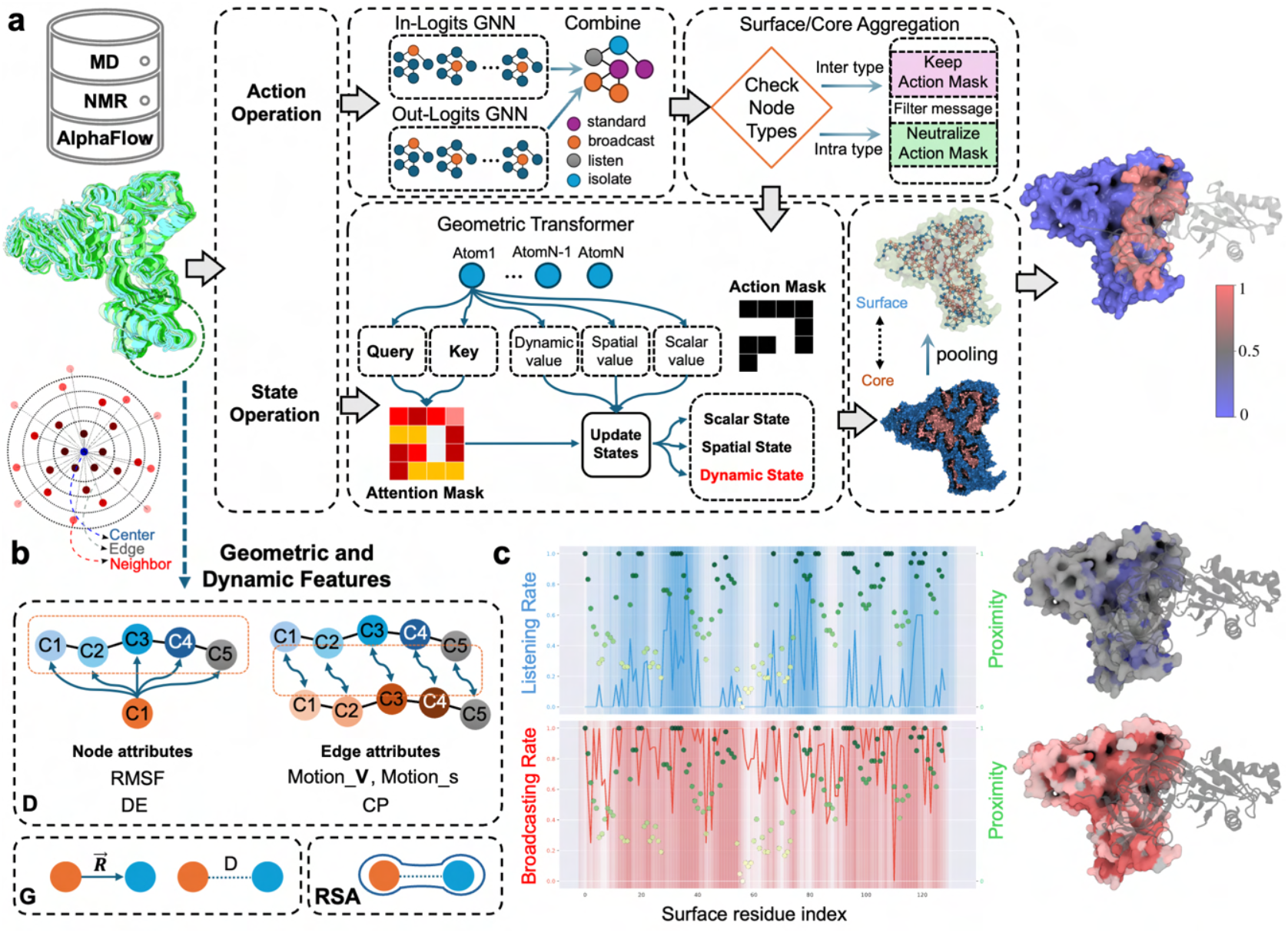
The computational workflow of DynamicGT. (**a**) DynamicGT processes the protein conformational ensemble derived from MD simulations, NMR, and AlphaFlow. The architecture employs cooperative GNNs, where the first two GNNs determine the actions of each atom, effectively constraining message passing between surface and core residues. With each layer of the architecture, the number of residues increases to emphasize closer neighbors. Subsequently, a geometric transformer updates the atomic states. The final block pools atomic information to residues and decodes this into predicted probabilities. (**b**) RMSF and DE are dynamic features associated with nodes, while motion V, motion s, and CP are defined for edges. RSA is calculated by averaging across multiple conformations. Additionally, displacement vectors and distances are defined to capture spatial and geometrical features of the protein structure. (**c**) Visualization of the broadcasting rate (red) and listening rate (blue) of surface residues, illustrating their relationship with the proximity of each residue to the interface.

We proposed a novel time-free feature extraction pipeline using vectors as features in a rotationally equivariant manner (Supplementary Note S1), which effectively integrates dynamic data. These features are detailed in the Methods section. In short, we proposed communication propensity (CP) which complements the geometric distance feature, and the motion vector (motion-V) and motion scalar (motion-s) that add dimensionality to edge attributes by capturing dynamic correlations between atoms. The relative accessible surface area (RSA) is computed by averaging across conformations to minimizes bias toward any single conformation and enhance core/surface node discrimination. Additionally, the root mean square fluctuation (RMSF) and the directional entropy (DE) characterize the flexibility and variability of the atomic movements (Figure 1b).

The underlying intuition behind cooperative GNNs is that the action of each node constrains its communication, which, in turn, affects the final outcomes. To investigate these interactions, we analyzed the in-logit (listening) and out-logit (broadcasting) actions across benchmark datasets, averaging them at the residue level. As a proof of concept, we visualized these embeddings for a subset of high-performing predictions (Figure 1c, Supplementary Figure S4a), where increased communications were observed around the binding sites. Moreover, comparing action states between surface and core residues revealed that surface residues exhibit a higher propensity to listen, whereas core residues predominantly broadcast (Supplementary Figure S4b).

In binding site prediction, integrating the distance of residues from the actual interface into the loss function during training enables the model to prioritize interface sites (as a patch of residues) over individual residues. This approach penalizes false positives proportionally to their distance from the binding interface, rather than uniformly treating all false positives as equals. For instance, a false positive residue located on the rim of a protein interface is less detrimental than one located at a distant site. To address this challenge, we proposed the GeoLoss function, an extension of the focal loss [41] designed to handle class imbalance in binary classification tasks. This spatially-aware loss function adjusts based on the distance of each residue to the interface, allowing the model to differentiate between core and surface residues. Specifically, distances for core residues are computed using the Euclidean metric, while surface residues are evaluated using a geodesic metric by computing the surface mesh. Both types of distances are normalized to ensure a uniform distribution, such that the residue farthest from the interface within the core has the same loss contribution as the farthest residue on the surface (Supplementary Figure S5). The GeoLoss can be formulated as follow:

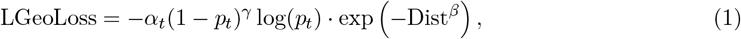

where *Dist* is the calculated distance of each residue from the interface, and *β* is a hyperparameter that controls the sensitivity of the distance (Figure 2).

**Figure 2:**
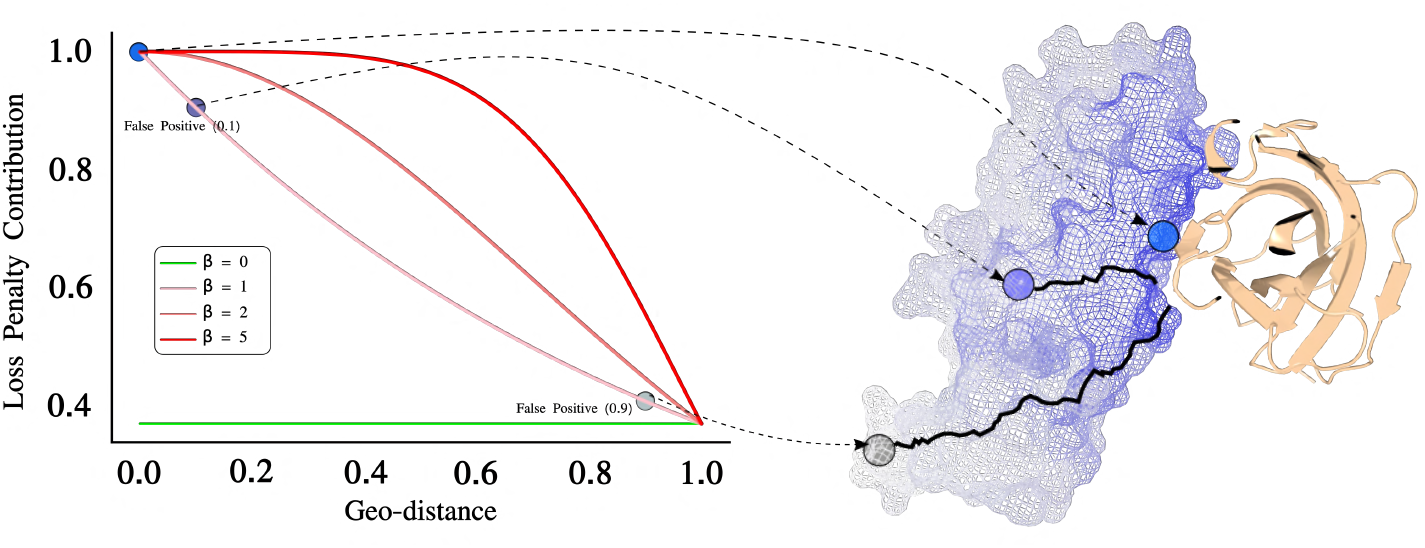
Spatially-aware loss penalty calculations in GeoLoss. Impact of *β* on GeoLoss penalty decay. Left: Loss penalty decreases with geodesic distance (Dist), shown for *β* = 0, 1, 2, 5. Right: Schematic mapping of false positives at distances 0.1 and 0.9 over the protein structure.

A primary challenge in training our model was the limited availability of dynamic data. To address this, we provided an extensive set of MD simulations and collected NMR complexes from PDB (Supplementary Figure S6). Considering the PDBbind PPI database [42], we curated a set of 1,115 protein chains in their bound forms (after redundancy and structural similarity removal) to carry out MD simulations (Supplementary Table S1, and Supplementary Figure S7). Performing three replicates of 500 microseconds for each entry, we obtained an overall 1-ms MD simulation data, and combined these data with other existing MD resources such as ATLAS (unbound/single chain form) [43], and MDDB (https://mddbr.eu/, see Methods section for more details), creating a comprehensive dataset of conformational ensemble for ∼ 3000 protein chains. While NMR entries do not fully encapsulate the dynamic conformational landscape of proteins in solution, they contain valuable information on conformational heterogeneity, particularly in regions of structural flexibility such as IDRs.

MD simulations as a gold standard provided a systematic probe over conformational states, increasing generalizability with varying RMSF ranges (Supplementary Figure S8). To further expand this training data and to facilitate running the model in the inference mode (as MD simulations are costly), we incorporated generated conformations by AlphaFlow to compensate the lack of data (Supplementary Figure S6), and designed a dynamic-aware (no AlphaFlow) and a heterogeneity-aware (MD, NMR, and AlphaFlow) model. The quality assessment was done using predicted local distance difference test (pLDDT) showing that 92 percent of data entries obtained a pLDDT greater than 70%, despite most of them containing disordered regions (Supplementary Figure S9). Finally, Considering both bound and unbound states allowed the model to make accurate predictions regardless of whether the input protein is in a bound or unbound conformation.

### Predicting binding regions with disordered binders

IDRs represent a significant subset of proteins characterized by the absence of a stable 3D structure in isolation. These regions can undergo disorder-to-order transitions upon binding to receptor proteins (i.e., folding upon binding) or remain intrinsically disordered even in the bound state. Due to their inherent conformational heterogeneity, accurately predicting binding interfaces on both IDRs and their receptor partners remains a major challenge in structural biology. Given that DynamicGT is designed to learn and capture dynamic heterogeneity in proteins, we evaluated its ability to predict binding regions on both IDRs and their receptor proteins. To ensure a comprehensive assessment, we considered the full spectrum of IDR binding modes, including disorder-to-disorder interactions, disorder-to-order transitions (folding upon binding), and mutual folding, where both interacting partners are disordered in isolation but adopt secondary structures upon binding (Figure 3a).

**Figure 3:**
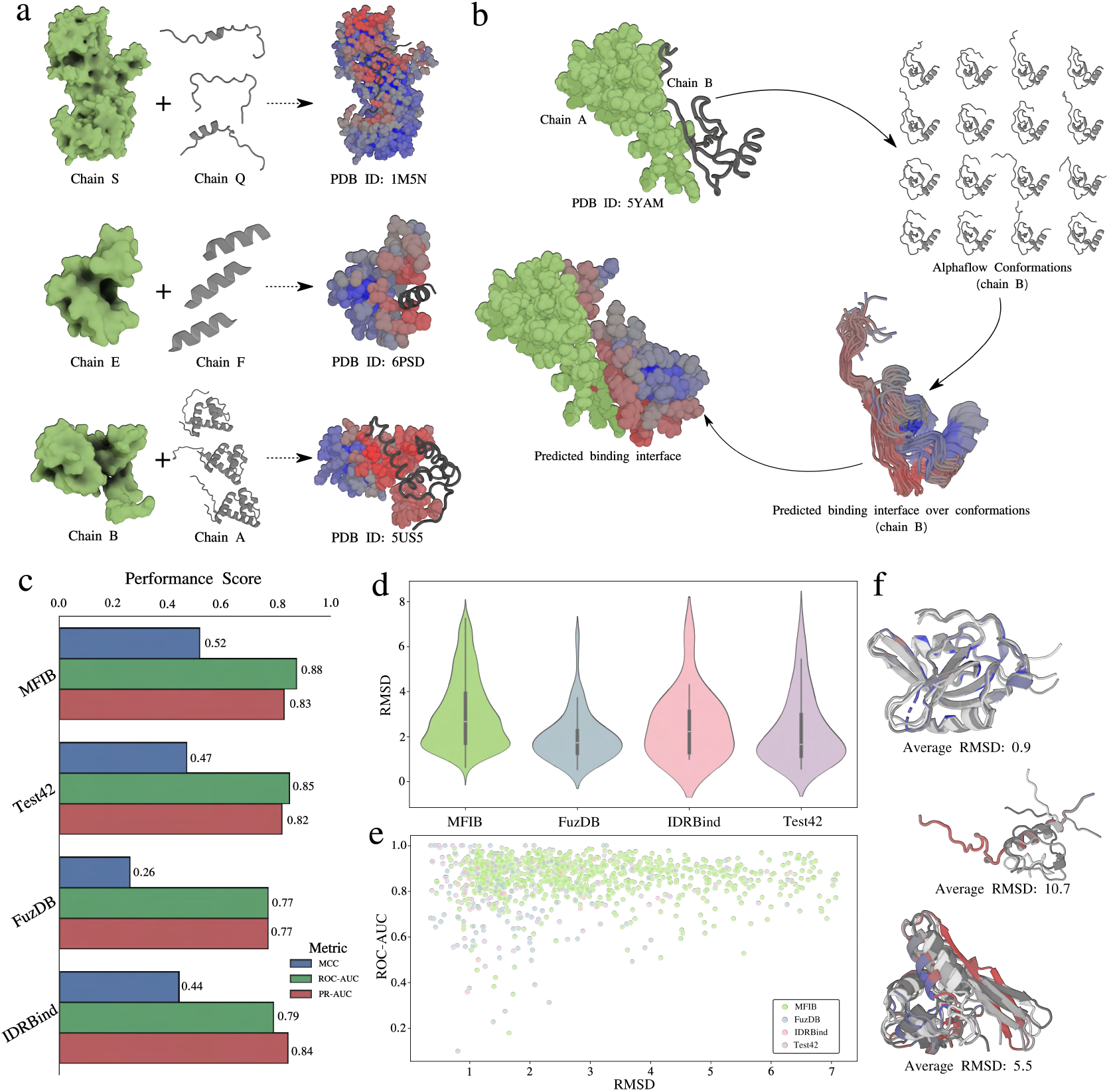
DynamicGT performance in IDR-mediated PPIs. **(a)** IDR mode of bindings, including disorder-to-disorder, folding upon binding, and mutual folding. The examples illustrated here (PDB IDs: 1M5N, 6PSD, 5US5) are derived from benchmark sets after model prediction, where receptor is depicted in green, and IDR in gray. **(b)** Workflow for predicting PPI probabilities based on conformational heterogeneity. The conformational ensemble for target chain (B) is generated via AlphaFlow and input into DynamicGT, yielding predicted interface probabilities. These probabilities are mapped over the conformations and evaluated against the native complex interaction (chain A). **(c)** Model performance on four benchmarks, MFIB, Test42, FuzDB, IDRBind evaluated based on MCC, ROC-AUC, and PR-AUC metrics. **(d)** RMSD distribution across all benchmarks (with outliers removed), showing the variation of generated ensembles with respect to the native structure. **(e)** RMSD vs ROC-AUC plot across all benchmarks (with outliers removed), to evaluate the impact of RMSD distribution over performance metrics. **(f)** Illustrative examples of conformations with low, medium, and high RMSD distributions.

To address these diverse binding patterns, we benchmarked DynamicGT across multiple datasets: FuzDB [22], a curated collection of protein–IDR interactions characterized by heterogeneous, fuzzy binding; the IDRBind dataset [15], which includes both bound and unbound protein–IDR complexes; Test42 [19], a rigorously compiled benchmark of 42 protein–peptide complexes with no overlap with the AlphaFold2 training set; and the MFIB benchmark [44], containing both homomeric and heteromeric IDR–IDR complexes that undergo mutual folding upon binding. DynamicGT was run in inference mode on conformational ensembles generated by AlphaFlow using single-chain inputs (sequences without structural templates), as illustrated in Figure 3b. To mitigate redundancy bias, we applied a cluster-based splitting strategy and excluded entries with sequence similarity to the training set (see the Methods section for details).

The model’s performance across all benchmarks is summarized in Figure 3c, highlighting metrics such as the Matthews correlation coefficient (MCC), median area under the receiver operating characteristic curve (ROC-AUC), and median area under the precision-recall curve (PR-AUC). Among all benchmarks, MFIB demonstrated the highest performance, achieving a median ROC-AUC of 0.88. In comparison, Test42, FuzDB, and IDRBind obtained ROC-AUC values of 0.84, 0.77, and 0.79, respectively. Notably, the strong performance on MFIB suggests that DynamicGT effectively captures the cooperative folding dynamics characteristic of mutual folding events, where dynamic features are likely more pronounced. Further analysis of the root mean square deviation (RMSD) distributions relative to reference static structures (Figure 3d) revealed larger conformational variations in MFIB entries, consistent with their dynamic folding behavior. Intriguingly, DynamicGT maintained consistent predictive performance (ROC-AUC) even for entries with substantially larger RMSD values (Figure 3e-f). This robustness suggests that the model’s dynamic feature representations are not overly sensitive to structural variability, reinforcing its potential to generalize across diverse conformational landscapes.

To dissect the model’s performance on structured versus flexible regions, we analyzed the secondary structure content of FuzDB and IDRBind entries. Residues were categorized into flexible regions (turns, bends, or unstructured segments) and structured regions (*α* helices, β strands). As expected, the model performed better on structured residues, achieving a ROC-AUC of 0.83 and an average precision of 0.51. Nevertheless, predictions on flexible regions remained robust, with a ROC-AUC of 0.78 and an average precision of 0.40 (Figure 4a). These results highlight DynamicGT’s ability to capture binding signals even within intrinsically disordered segments, although with reduced precision compared to structured regions.

**Figure 4:**
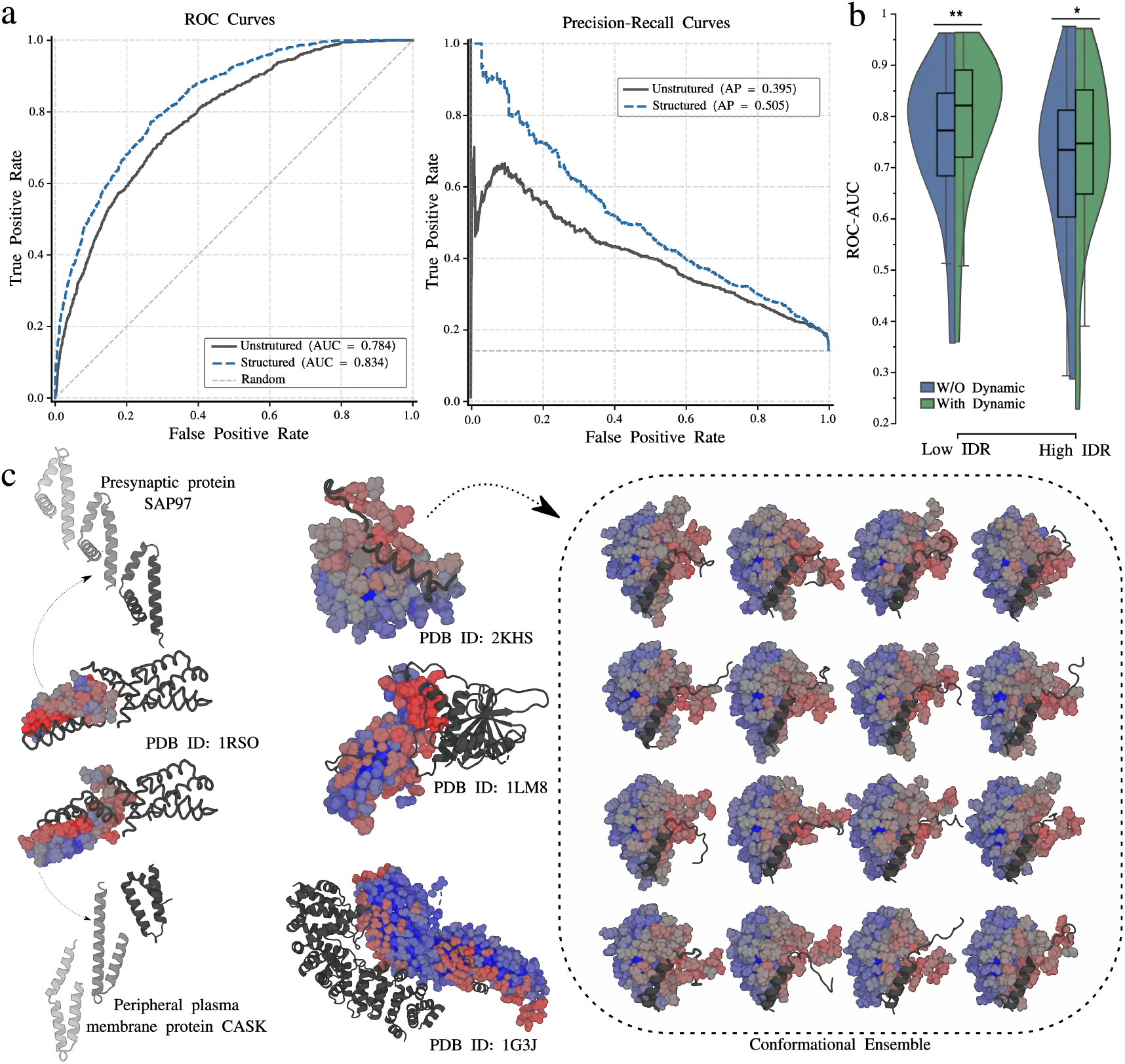
Evaluating the impact of dynamic features on structured and IDR-rich interfaces. (a) Residue-wise performance comparison for structured (ROC-AUC = 0.834, PR-AUC = 0.505) and unstructured (ROC-AUC = 0.784, PR-AUC = 0.395) regions using ROC-AUC and PR-AUC metrics. **(b)** ROC-AUC performance of models with (green) and without (blue) dynamic features across proteins with low and high extent of IDRs. Dynamic features significantly improved performance (\**p ≤* 0.05, \*\**p ≤* 0.01). **(c-left)** Structure of PDB ID: 1RSO with a stoichiometry of A_2_B_2_; a MFIB example. Conformational ensembles for each chain were generated using AlphaFlow, and interfaces were predicted using DynamicGT. **(c-right)** Examples of model performance evaluated with respect to the native PPIs. The left panel shows structures from PDB IDs 1G3J, 2KHS, and 1LM8. The right panel displays the entire conformational ensemble predicted for the 2KHS (Nase121:SNase(111-143) complex).

To further evaluate the impact of IDRs on model performance, we computed per-residue disorder probabilities using MetaPredict [45]. Residues were classified as ordered or disordered based on a probability cutoff of 0.5. We then compared the performance of DynamicGT against a no-dynamic baseline model (by ablating dynamic features) across two groups: proteins with a low extent of IDRs and those with high IDR content (i.e., disordered regions above median). In both groups, DynamicGT significantly outperformed the no-dynamic model, underscoring the relevance of dynamic features. However, the performance gap between DynamicGT and the baseline was sharper for proteins with low IDR content (Figure 4b). This can stem from challenges associated with sampling the vast conformational space of highly disordered proteins and ineffective distinction of surface and core residues (no core residue can typically be considered for highly disordered proteins) which reduce the impact of cooperative message-passing.

Representative predictions are showcased in Figure 4c (more examples in Supplementary Figure S10), including complexes with multiple binding interfaces and an MFIB heteromeric complex. These examples illustrate the model’s ability to capture binding regions across diverse conformational states of each binding partner. Notably, even in cases where binding involves transient or partially ordered intermediates, DynamicGT effectively identified key interface residues, highlighting its potential for predicting dynamic, context-dependent binding events.

### From Highly Flexible to Transient and Static Interactions

PPIs can be broadly categorized as transient or permanent [46], with transient interactions posing greater challenges for predictive models due to their dynamic and often weakly stable nature. While DynamicGT is designed to enhance performance on flexible and disordered regions, we sought to further evaluate its effectiveness in predicting both transient and static interactions, where state-of-the-art models have shown strong performance [13, 36, 37, 39]. To ensure direct comparability with existing methods, we assessed our model using two widely recognized benchmarks: the “Testing Transient” dataset, comprising 53 protein–protein complexes [37], and the “Test60” benchmark [36]. To prevent data leakage and ensure unbiased evaluation, a cluster-based dataset splitting strategy was employed (see Methods for details).

Since many static models rely on bound structures to predict binding interfaces, potential label leakage—where the bound conformation implicitly contains binding information—could influence the performance. To address this, we performed two distinct evaluations of DynamicGT on the “Testing Transient” benchmark. The first model, trained exclusively on MD simulation data, was tested on an MD-derived benchmark constructed from bound structures, achieving a median ROC-AUC of 0.868, with marginal improvements over leading static models. The second, a generalized model trained on both MD simulation and AlphaFlow-generated data, was evaluated on conformational ensembles generated from single-sequence inputs without structural templates. This model also achieved a comparable ROC-AUC of 0.851, highlighting its robustness even in the absence of explicit structural information (Figure 5a, and Supplementary Table S2). We further evaluated DynamicGT on the “Test60” benchmark and compared its performance against several state-of-the-art models, including EquiPPIS [36], GraphPPIS [47], MaSIF-site [37], and SPIDER [48]. DynamicGT demonstrated superior performance, achieving the highest PR-AUC of 0.479 (Figure 5a).

**Figure 5:**
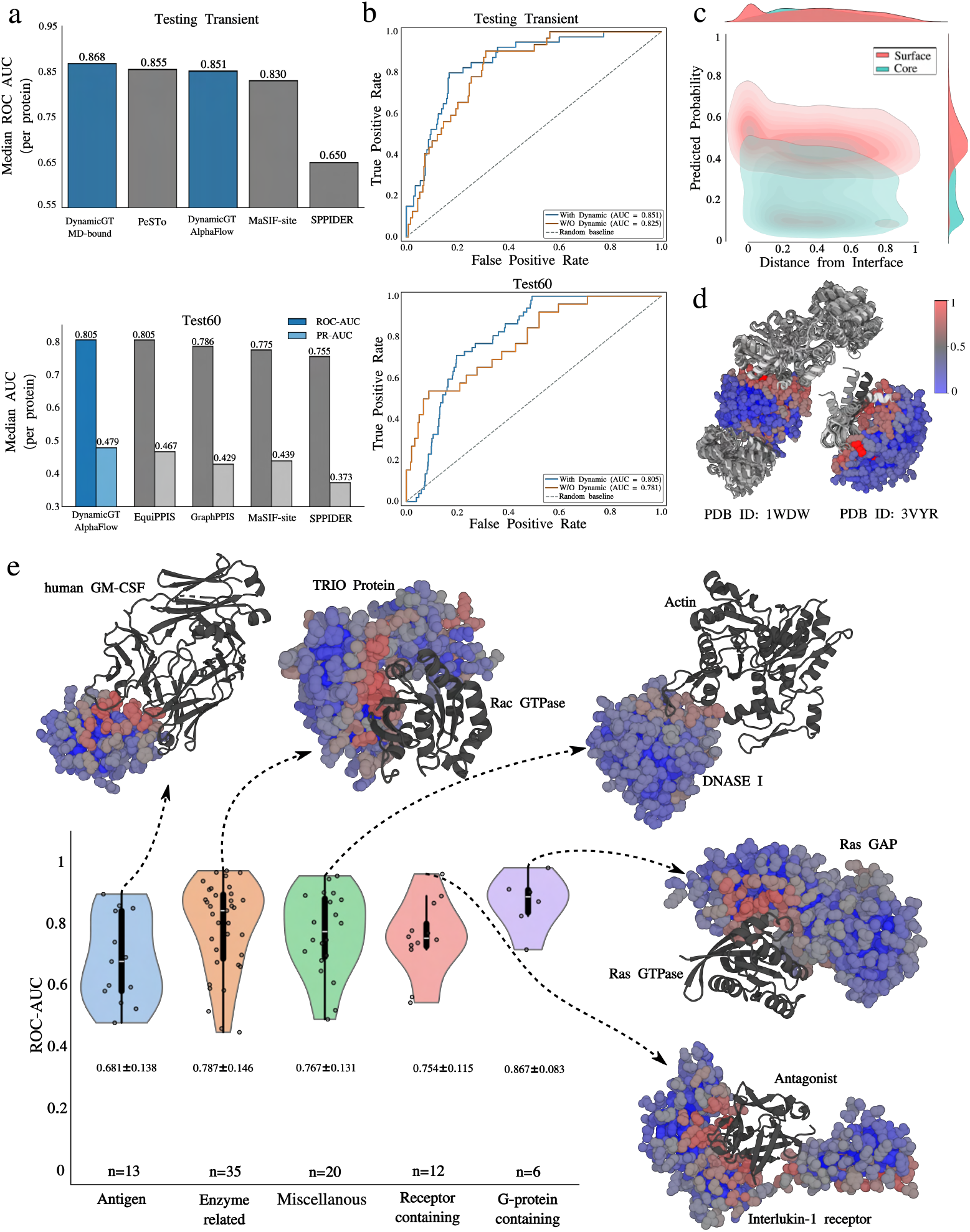
Prediction of binding sites in static and transient PPIs. (**a**) Comparison of DynamicGT model with the state-of-the-art site prediction approaches including PeSTo, MaSIF-site, and SPPIDER over two benchmark sets of single-chain “Testing Transient” and “Test60”. The performance of DynamicGT model trained over only MD simulations data is shown for comparison as well. (**b**) PRC curves representing the performance of DynamicGT on “Testing Transient” and “Test60” benchmarks, where the network is trained with only geometric (no dynamic), and geometric plus dynamic features. (**c**) Predicted probability distribution of residues being at the protein interface versus core against the distance from the interface, across the entire of “Testing Transient” benchmark. The contour and marginal density plots highlight the separation between interface (cyan) and core (red) residues. (**d**) Two test examples of DynamicGT predictions showing well-formed binding sites having conformational states of the input protein. (**e**) Performance of the model on the ZDock benchmark, shown with structural examples. The number of entries, ROC-AUC, and standard deviation are indicated.

To quantify the contribution of dynamic features, we conducted a series of ablation studies by systematically removing key components, including dynamic features (RMSF, motion), cooperative message passing, and AlphaFlow-generated conformations. We first compared DynamicGT (heterogeneity-aware model) with a no-dynamic baseline model (retaining only cooperative message passing and GeoLoss) across the “Testing Transient” and “Test60” benchmarks. The dynamic-enriched model consistently outperformed the static model, achieving 2.6% and 2.4% higher ROC-AUC scores on the “Testing Transient” and “Test60” benchmarks, respectively (Figure 5b). These results highlight the added value of incorporating dynamic information, even when predicting static or transient binding interfaces.

Next, we assessed the importance of individual features and cooperative message passing using the dynamic-aware model. Ablating dynamic features resulted in a 3% drop in ROC-AUC compared to the full dynamic-aware model (Supplementary Figure S11a). To further evaluate cooperative message passing, we disrupted the RSA feature by introducing noise, effectively removing the distinction between core and surface residues. This modification led to the lowest ROC-AUC among all tested configurations, underscoring the critical role of cooperative message passing.

Finally, in the heterogeneity-aware model, the impact of dynamic features was less pronounced, but the model incorporating them still achieved the best overall performance (Supplementary Figure S11b). These findings suggest that while heterogeneity-aware predictions benefit from broader conformational sampling, explicit dynamic information derived from MD simulations remains a key factor for interface predictions.

The impact of GeoLoss is also illustrated in Figure 5c. Training the model with GeoLoss on the transient benchmark revealed three distinct residue probability distributions: (i) core residues consistently exhibited low predicted probabilities regardless of their proximity to the interface, (ii) surface residues near the binding interface displayed high probabilities, and (iii) surface residues farther from the interface showed moderate probabilities. This distribution enables the model to tolerate close false positives by recognizing them as potential rim residues—those peripheral to the core interface—while effectively distinguishing residues based on structural depth and interaction likelihood, improving the model’s spatial awareness of interaction sites. Representative predictions (Figure 5d and Supplementary Figure S12) also illustrate the model’s ability to accurately identify multiple binding interfaces within a single protein chain.

To further evaluate DynamicGT in specific application contexts, we conducted additional assessments using the ZDock benchmark [49], which includes both bound and unbound protein structures. We excluded entries with sequence similarity to the training set and removed all antibody chains, as their well-defined binding sites could artificially influence performance metrics. The remaining entries were categorized based on complex type and used as targets for interface prediction (Figure 5e, Supplementary Figure S13). Among these, chains from GPCR complexes exhibited the highest performance, achieving a median ROC-AUC of 0.89, highlighting the model’s robustness in capturing the dynamic conformational landscapes characteristic of membrane-associated receptors. In contrast, antigen–epitope mapping posed the greatest challenge, with a median ROC-AUC of 0.68, likely due to the structural plasticity and high sequence variability of antigenic epitopes. For enzyme-related complexes, receptor-containing assemblies, and miscellaneous protein interactions, the model achieved median ROC-AUC values of 0.84, 0.75, and 0.77, respectively. These results showed DynamicGT’s adaptability across a wide range of interaction types while identifying areas—such as antigen–epitope recognition—where incorporating additional dynamic features or specialized training strategies may further improve predictive performance.

## Discussion

In this study, we introduced DynamicGT, a novel deep learning framework that leverages advanced GNN architectures with cooperative message-passing to capture the dynamic heterogeneity of proteins. DynamicGT demonstrates the ability to extract meaningful dynamic features without explicitly modeling trajectories as time-series data. We showed that incorporating these dynamic features significantly enhances binding interface prediction, particularly in systems where conformational flexibility is critical, such as IDRs. By training on large-scale MD simulation data—considered the gold standard for probing protein dynamics—we developed a dynamic-aware model capable of robust interface prediction. Additionally, we extended this approach by integrating predictive conformational ensembles generated by AlphaFlow and NMR-derived structures from the PDB, further enhancing the model’s sensitivity to structural heterogeneity, and facilitate running the model on inference mode for both benchmarking and end users.

While graph transformers have emerged as powerful tools for binding interface prediction [13, 39], surface-based models have highlighted the critical role of surface residues, often at the expense of oversimplifying the structural complexity of proteins [37]. DynamicGT bridges this gap through cooperative message-passing that integrates information from both surface and core residues. By incorporating dynamic features, our model learns hidden structural patterns beyond static representations, offering a more comprehensive view of protein interaction landscapes.

We performed extensive evaluations of DynamicGT across diverse benchmarks, including a wide range of PPI types and dynamic contexts. Our results demonstrate substantial improvements in predicting binding sites for IDRs and their receptor proteins. Remarkably, DynamicGT achieves performance on par with state-of-the-art static predictors, despite being trained on significantly less data. Moreover, while many existing methods specialize in either IDRs or structured receptor proteins, DynamicGT maintains robust performance across both categories. Importantly, the model performs consistently well on both bound and unbound structures, as evidenced by its performance on the ZDock benchmark, which includes a diverse set of protein complexes with varying functional roles.

DynamicGT holds considerable promise for advancing the prediction of protein complexes involving flexible regions or IDRs. Its architecture can be seamlessly integrated with end-to-end structure prediction models such as AlphaFold or RoseTTAFold [50], potentially enhancing their accuracy in disordered and flexible regions. Future directions include fine-tuning DynamicGT for partner-specific interactions and extending its application to other biomolecular interaction types, such as protein–ligand and protein–NA complexes. Additionally, by learning dynamic features through cooperative message-passing, DynamicGT opens new avenues for inferring hidden structural relationships. These insights can be further explored to address complex biological questions, such as mutational landscape or stability of proteins.

## Data availability

The training data for this study, including clustered representatives of MD simulations, and all AlphaFlow generated models are available at Zenodo through https://doi.org/10.5281/zenodo.14833854.

## Code availability

The code, model parameters, details regarding the training, test data split, and benchmarking are available at GitHub address https://github.com/amisteromid/DynamicGT.

## Acknowledgment

This work was granted access to the HPC resources of IDRIS under the allocation 2024-GC010715460 (granted to YK and HK) made by GENCI. HK was supported by the French Agence Nationale de la Recherche (ANR), under grants ANR-22-CPJ2-0075-01, and ANR-24-CE45-4243-01.

## Contributions

OM, YK, and HK conceived the idea. YK performed all MD simulations and data analysis. OM developed the deep learning model in discussion with SG and HK, and performed the data analysis. HK and YK supervised the project. OM and HK wrote the initial draft with input from all authors.

## Methods

### Setup of MD simulations

For every system, we performed three replicates of 500 ns MD simulations starting from the experimental structure. First we removed all the ligands, but kept the ions. Then, missing residues were added using MODELLER for consecutive gaps of maximum 5 residues or with AlphFold3 is the gap was longer than 5 amino acids. For the AlphaFold3 predictions, we chose the one with the highest pLDDT score. MD simulations were carried out with the GROMACS 2024.2 [51] using the CHARMM36m force field parameter set: (i) Na^+^, Cl ^−^ counter-ions were added to reproduce physiological salt concentration (150 mM solution of sodium chloride), (ii) the solute was hydrated with a dodecahedron box of explicit TIP3P water molecules with a buffering distance of up to 12 Å, and (iii) hydrogen atoms were added and the environment of the histidine was checked using the Reduce software [52]. For each system, the energy minimization was performed by steepest descent algorithm for 5000 steps, to minimize any steric overlap between system components. This was followed by an equilibration simulation in an NPT ensemble at 310 K, allowing the lipid and solvent components to relax around the restrained protein. All the protein and lipid non-hydrogen atoms were harmonically restrained, with the constraints gradually reduced in 6 distinct steps with a total of 0.375 ns. During the system equilibration steps, the pressure was maintained at 1 bar with Berendsen thermostat and barostat [53], respectively. For every system, three replicates of 500 ns, with different initial velocities were performed in the NPT ensemble using the time step of 2.0 fs. The temperature was kept constant at 310 K using V-rescale thermostat [54] and a constant pressure of 1 atm was maintained with C-rescale barostat [55]. Isotropic pressure treatment was applied to the system. The bonds involving hydrogen atoms were constrained using the LINCS algorithm [56], while the electrostatic interactions were calculated using Particle Mesh Ewald method [57], and the coordinates of the system were written every 100 ps.

### Training data generation

We utilized two pre-existing resources of MD simulations data: the MMB node of the MDDB databank (https://mmb.mddbr.eu/), which contains 83 entries, and the Atlas of Protein Molecular Dynamics (ATLAS) [43], a database of single-chain MD simulations. For the ATLAS dataset, we excluded chains classified as monomers in the PDB, reducing the selection from 1,390 to 673 chains. To further extend our dataset, we performed MD simulations across several datasets. First, we utilized PPIs from the PDBbind database [42]. Second, we curated single-chain antigens from the SAbDab database [58], prioritizing entries with both heavy and light chains, as well as those annotated with binding affinity. For both datasets, structural redundancy was eliminated using MM-Align [59], ensuring a diverse and non-redundant selection. Finally, for each entry with available MD simulations, we performed RMSD-based clustering of the snapshots dynamically by varying the cutoff. This approach allowed us to generate a certain number of representative structures. For AlphaFlow-generated conformations, we used the distilled version, obtaining 25 conformations per chain, resulting in a total of around 13,000 chains. For the training dataset, conformations were filtered by comparing AlphaFlow-generated structures to their native counterparts, retaining only those with a RMSD below 4Å. In addition, all NMR solution PPIs from the PDB were used.

### Alignment strategy

To ensure uniformity across the dataset, the conformations of each PDB structure—whether derived from MD simulations, NMR, or AlphaFlow—were re-aligned using the Kabsch algorithm [60] via `superimposer` in Biopython before extracting features.

### Benchmark sets

We performed MD simulations on complexes from the “testing transient” dataset and conducted extensive benchmarking using AlphaFlow. The benchmarks included “testing transient”, “test60” [36], FuzDB [22], IDRBind [15], MFIB [44], Zdock [49], and a set of 42 protein-peptide complexes [19]. To evaluate model performance on interactions involving intrinsically disordered regions (IDRs), we utilized FuzDB, IDRBind, MFIB, and the 42 protein-peptide complex dataset. The “set-42” dataset comprises proteins with IDR-containing binding partners, demonstrating the limited performance of AlphaFold-Multimer in such cases. The Mutual Folding Induced by Binding (MFIB) database includes 1,210 complexes exclusively formed by intrinsically disordered proteins (IDPs). FuzDB compiles complexes in which intrinsic disorder is maintained upon interaction with a partner, while IDRBind contains complexes where at least one partner has experimental evidence of being intrinsically disordered prior to binding. The “testing-transient” and “test60” datasets are widely used for general benchmarking and comparison with state-of-the-art methods. Zdock provides a diverse dataset of proteins in both bound and unbound states for evaluating protein-protein docking models, including categorization based on interaction types (rigid-body and medium difficulty) as well as functional classifications.

### Inference

For model inference, the conformational ensemble of a given protein is required. This can be obtained either by utilizing AlphaFlow or through MD simulations, with the latter providing more accurate dynamic features but computationally demanding.

### Feature extraction strategy and labeling

Feature extraction was performed at the individual chain level, with chains separated for complexes subjected to MD simulations. Binding site residues were annotated from MD representative structures in the case of complexes, capturing the full range of sampled conformations. For single-chain simulations, interface residues were determined from static structural data.

### Ground truth labels

We used a SASA-based method to define interface residues by finding those that are buried in complex form and exposed in isolate form as seen previous studies [48, 61]. We used a cutoff of a 4% change in SASA, and this is applied to each conformation to avoid missing transient interactions. Additionally, bound structures come with labels derived from all conformational states, ensuring that interface residues are comprehensively annotated across the dataset.

### Dataset splitting strategy

To remove redundancy, and handle data limitation at the same time, we used MMseqs2 [62] with a lenient threshold of 0.9. Then to avoid information leakage between the training, validation, and test splits, we split the data based on clusters made by 0.4 cutoff, rather than individual sequences. Additionally, clusters containing benchmark entries were entirely excluded from the training set.

### Feature definition and calculation

Each protein is represented as a graph where nodes denote atoms and edges represent atom-atom interactions. One-hot encoding of node types is done for 29 frequent elements found in PDB structures, plus 1 for rare elements. The node level features include: RMSF, and DE, and the edge level features include: CP, motion-V, and motion-s.

### Distance and displacement vector

We averaged the positions of each node across all conformations of a protein to calculate the pairwise displacement vectors between all nodes for that protein. These vectors were then normalized by dividing by their norms. Following the same protocol as the displacement vector, we calculated the pairwise distances between nodes to capture the magnitude of displacement vectors.

### Communication propensity (CP)

After computing the pairwise Euclidean distances between elements across different conformations, we calculate the variance of these distances relative to their mean as reported in [63],

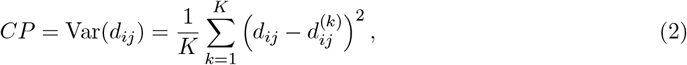

where 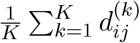 is the mean distance across all K conformations.

### Root mean square fluctuation (RMSF)

RMSF quantifies the average deviation of atoms from their mean positions, indicating flexibility within a conformational ensemble

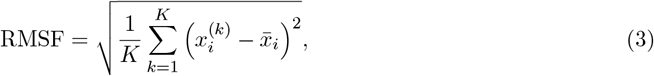

where 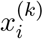 is the position of an atom in conformation *k* and 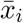 is the mean position of atom *i* over all conformations.

### Directional entropy (DE)

While RMSF focuses on both magnitude and direction, we extracted an additional feature to quantify the variance in the direction of these fluctuations. We stacked the normalized movement vectors for all conformations of each node, resulting in a tensor of size (number of conformations, number of nodes, 3) and used singular value decomposition (SVD) to extract singular values. First, we compute movement vectors 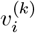, and normalize them

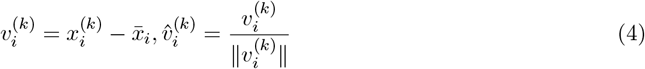

where 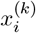 is the position of atom *i* in conformation *k*. Then, we stack these movement vectors 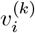 for all *K* conformations, and we perform singular value decomposition:

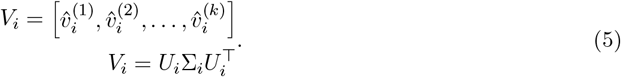

The three singular values obtained were normalized such that their sum equals 1. Finally, we calculated their directional entropy, which quantifies the spread or uniformity of the distribution of directions:

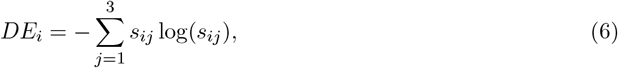

where

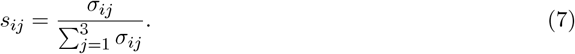

### Dynamic correlation

To capture the dynamic behavior of protein structures, we define a movement vector for each node in each conformation. This vector represents the displacement from the mean position of the node, averaged across all conformations, to its position in a specific conformation. By calculating the pairwise cross product of these movement vectors for each conformation, we generate a set of vectors for every pair of nodes, quantifying their dynamic correlation. We further define two features to describe this dynamic relationship:

#### Motion-V

The averaged cross-motion vector between each pair of nodes across all conformations is given by:

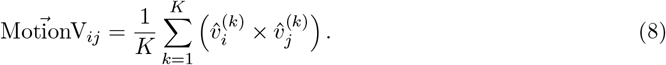

where 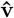 is defined as the movement vector.

Since movement vectors 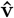 are normalized, the magnitude of MotionV reflects the degree of correlation, where 0 indicates perfect correlation and 1 represents minimal correlation.

#### Motion-S

The cosine similarity of these correlation vectors for each node pair, capturing the fluctuation in their relative motions, is defined as:

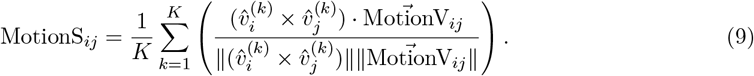

### Relative accessible surface area

The relative accessible surface area (RSA) measures the surface area of an amino acid residue that is accessible to a solvent, expressed relative to a standard or maximum value for that residue. We calculated the RSA for each conformation in the unbound state using the DSSP module from the Biopython package [64]. A probe radius of 1.4Å was employed, along with the default parameters provided by DSSP. The RSA values were averaged for each residue across all conformations.

### Geometric and geodesic distance calculation

To calculate geometric distances, a hybrid approach is used that integrates surface mesh with structural information. The mesh calculation was done using MSMS [65]. The surface mesh is treated as a graph, where nodes correspond to surface vertices, and edges are defined by the triangular faces connecting them. Each C-alpha atom is assigned to its nearest surface vertex considering a threshold of 6Å. Distance calculations depend on whether a residue is on the surface or in the core:

#### Surface residues

Residues with mapped surface vertices have their distances computed using the shortest path along the surface mesh to the nearest interface residue.

#### Core residues

Residues without nearby surface vertices have their distances measured as the minimum Euclidean distance to an interface residue.

To ensure comparability, we normalize distances separately for core and surface residues. Core distances are normalized relative to the maximum observed core-interface distance, while surface distances are scaled based on the minimum and maximum path lengths along the surface mesh.

### Architecture

Inspired by the cooperative-GNN architecture [40] and building on state-of-the-art static geometric transformer architecture from PeSTo [13], our architecture integrates the geometric transformer with the power of surface-centric methods [37]. At its core, it consists of two tightly coupled modules for updating action and state of nodes that iteratively refine node states across 35 layers (Supplementary Figures S1–S3). Every five layers, the receptive field expands dynamically—nearest neighbors grow from 12 to 48—allowing the model to progressively encode both local and global structural context. As the architecture leverages conformational ensembles, we extracted both scalar and vector dynamic features. To provide more informative representations, atomic dynamic correlations are captured using the cross product rather than the inner product. We further demonstrate that this approach preserves both rotation equivariance and translation invariance (see Supplementary Note 1).

### Action operation

We employ two distinct GNNs to compute the in-logits and out-logits for each node. The in-logits dictate whether a node can receive information, while the out-logits determine if a node can transmit information. This setup results in four possible actions for each node that would be encoded in a mask matrix: standard (both receive and transmit), listen (only receive), broadcast (only transmit), and isolate (neither receive nor transmit).

### Surface/core aggregation

Nodes are categorized as either surface or core based on their RSA in the unbound state, using a threshold of 0.16 [66] to distinguish between buried (core) and exposed (surface) residues. A second mask matrix is created at this stage, distinguishing between inter-type (core-surface) and intra-type (core-core or surface-surface) node interactions. Finally, this matrix is merged with the action mask matrix so that the action constraints are only applied to inter-type interactions.

### State operation

Each node is assigned three states: scalar state, spatial state, and dynamic state, which are updated layer by layer using a geometric transformer designed to be equivariant to both rotations and translations.

### Pool layer

An attention mechanism is employed to pool atom-level embeddings into residue-level representations, preparing them for the decoding layer and final predicted probability.

### Computation time

The dynamic-aware model was trained on a single NVIDIA A40 GPU (45 GB) for 28 epochs, completing in 28 hours. Early stopping with a patience of 5 epochs was used to prevent overfitting. Hyperparameter optimization was conducted via grid search on a subset of the dataset, yielding a final architecture consisting of 20,907,456 trainable parameters. A primary bottleneck during dataset preparation was the calculation of distance-to-interface metrics for the loss function; however, this step is not required during inference.

## Supplementary Information

### Supplementary Notes 1

#### Proof of Rotation Equivariance and Translation Invariance for the vectorial Motion feature

Let 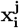 denote the protein coordinate of node *i* in conformation *j* for 1 ≤ *j ≤ K*, and let **R** be a 3 × 3 rotation matrix. The motion vector **motionV** is constructed as follows:

- The mean position is given by:

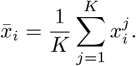
- Under rotation, 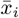 transforms as:

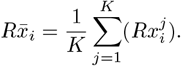
- The displacements 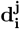 are given by:

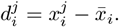

Under rotation, 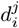 transforms as:

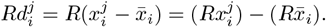
- The normalized displacements 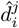 are given by:

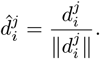

Under rotation, 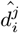 transforms as:

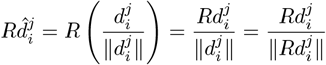

since ‖*Rd* ‖ = ‖*d* ‖ for rotation matrices.

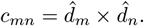
- The cross products *c*_*mn*_ for each pair (*m, n*) are given by: Under rotation, *c*_*mn*_ transforms as:

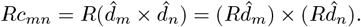
- The final motion vector **motionV** is given by:

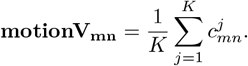

Under rotation, **motionV**_**mn**_ transforms as:

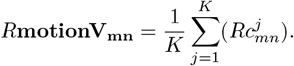

Thus, **motionV** is rotation-equivariant.

### Node and Edge Inputs

#### Node and edge feature packing

All scalar elements are utilized, including the dot product of *motionV* and *p*_motion_, as well as the norm of *p*_motion_. Consequently, *X*_*e*_ and *X*_*n*_ remain fully invariant under rotation.

### Multi-head Key, Query, Value Projection

- Generating keys (K), values (V), and queries (Q) from the edge and node feature packages using MLPs.
- Since inputs are rotation invariant, output of MLPs are fixed-size vectors (invariant).

### Vector Values Packing

1. The element-wise multiplication of the generated value with motionV. Value is invariant, so we would have *V* * *motionV*_nn_ *æ R*(*V* * *motionV*_nn_).
2. *p*_motion_ of the center node. 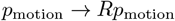.
3. *p*_motion_ of the neighboring nodes. 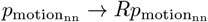.

Finally, we would have 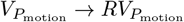.

### Attention Mechanism

- *Q* (from *nqm*) is invariant. *K*_*p*_ is derived from invariant *X*_*e*_, suggesting it’s a **fixed** descriptor, not a spatial vector.
- *M*_*p*_ = softmax(*QK*_*p*_) is invariant (softmax over 3*n* doesn’t alter transformation properties).
- 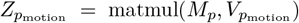. Since 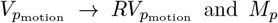 is invariant, we conclude that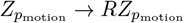.

### Projection (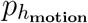)

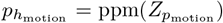, where ppm is a linear layer over [*Nh × S*] without bias: For each spatial dimension,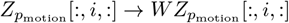 (same *W* size [Ns, Nh*Ns] applied independently). Since 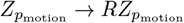, we get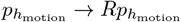.

Given that **motionV** is rotation-equivariant, we would have:

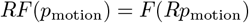

## Supplementary Figures

**Figure S1:**
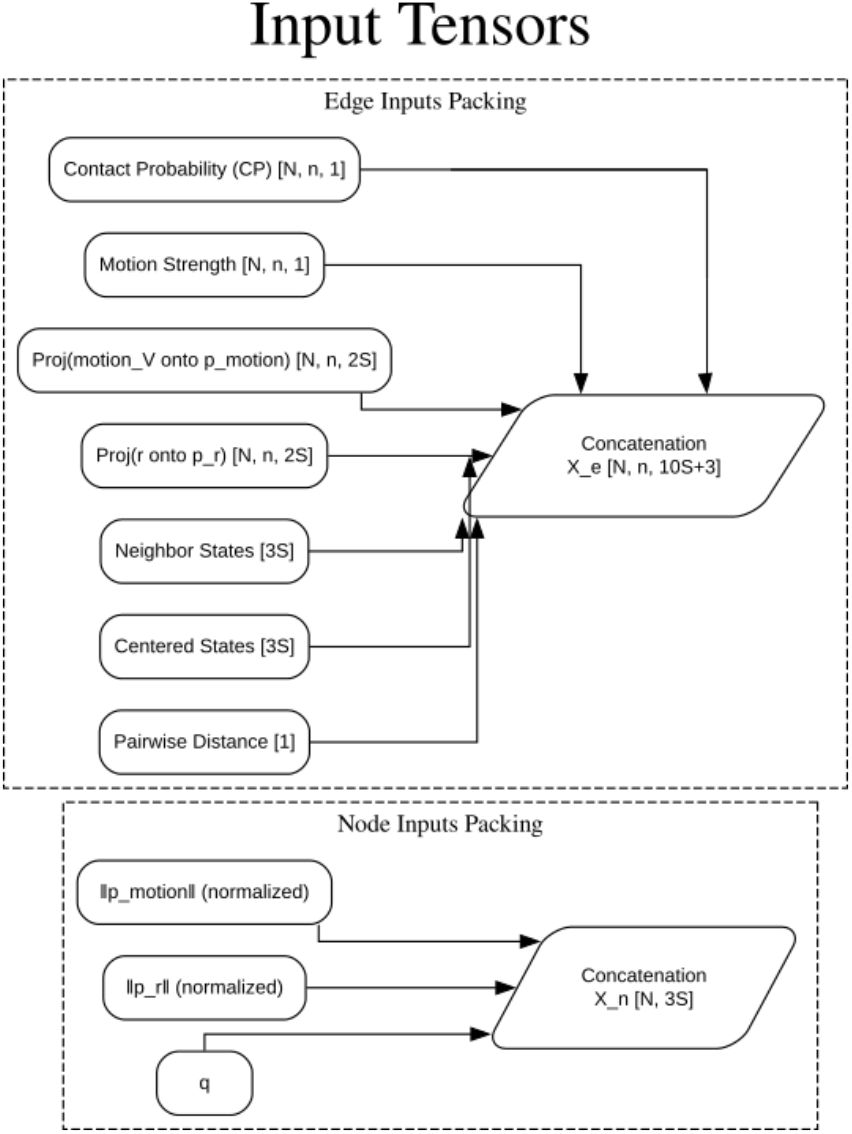
Node and edge feature packing. Diagram showing how features are used in the model. The node input q is a concatenation of atom-type, RMSF, DE, and RSA features. For the edge feature packing, each of the projections are calculated for the state of centered node and its neighbors.

**Figure S2:**
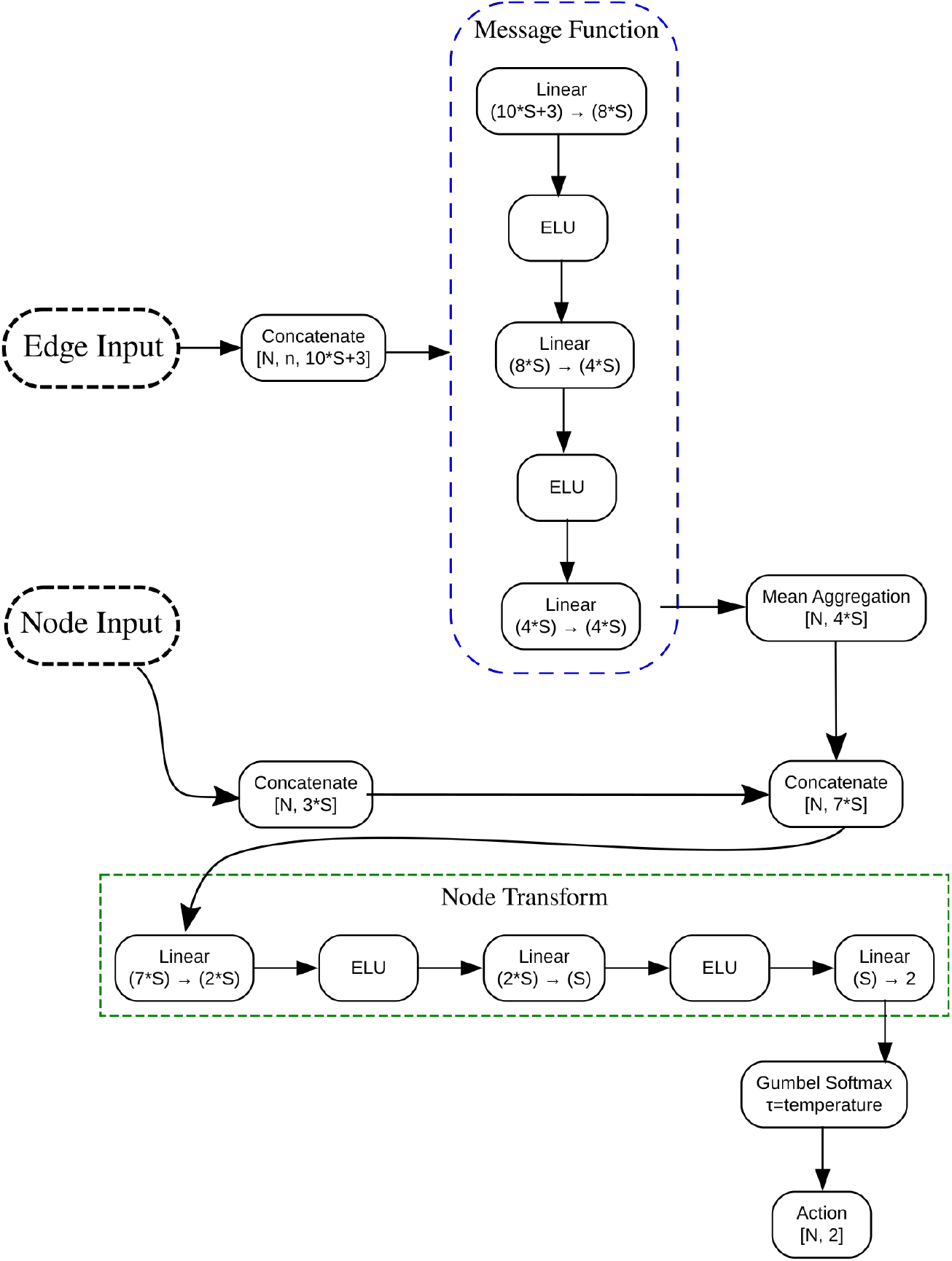
Complete architecture of the action update neural network module. Edge features are processed to generate messages, which are then aggregated at each node. The aggregated message is concatenated with node features to form a unified Representative, which undergoes a final transformation and discrete activation via Gumbel-Softmax.

**Figure S3:**
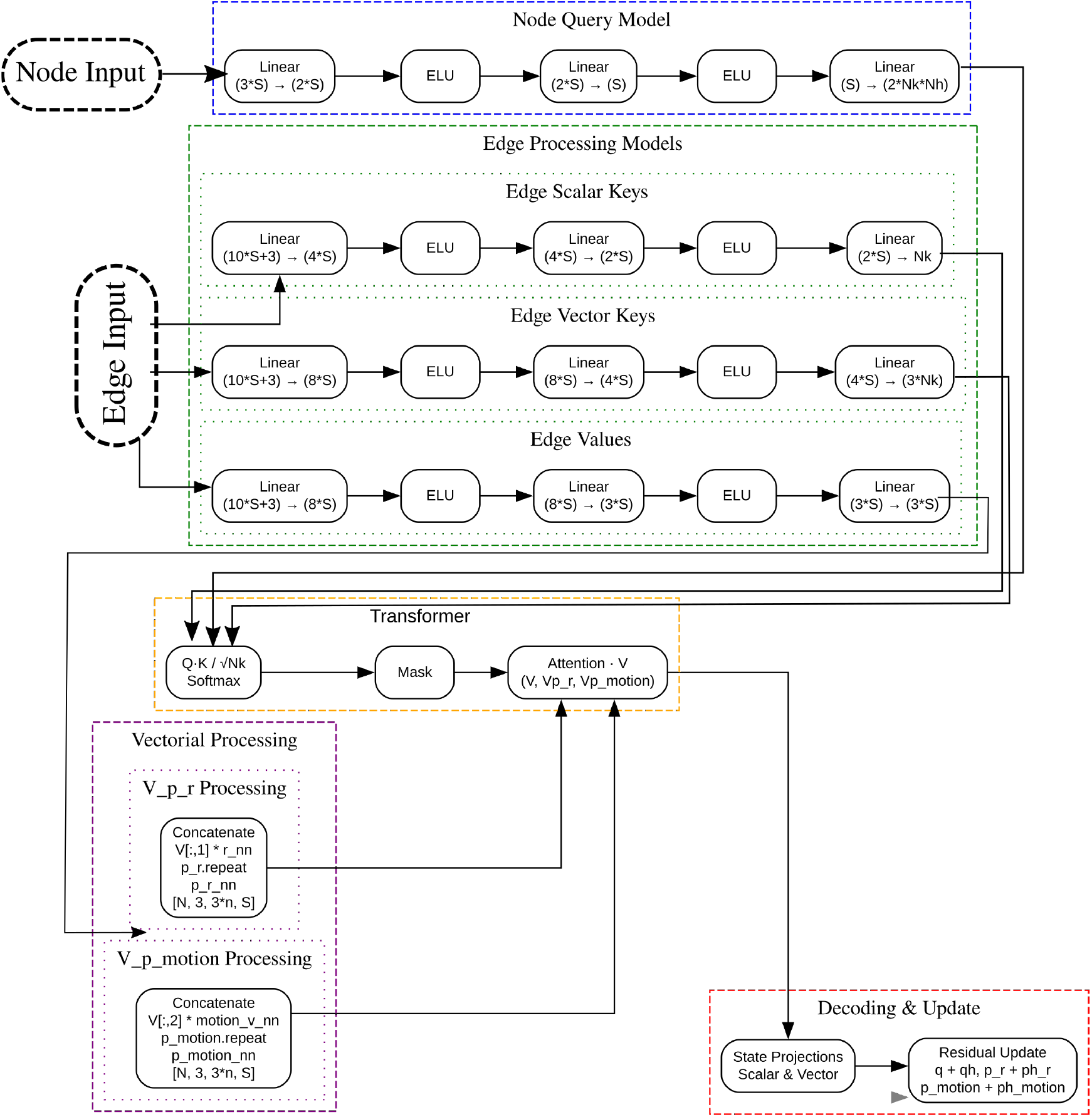
Complete architecture of the state update neural network module. Diagram depicting the state update operation, transformer module (inspired from PeSTo), followed by final decoding layer. The updated state is computed and summed with the previous state.

**Figure S4:**
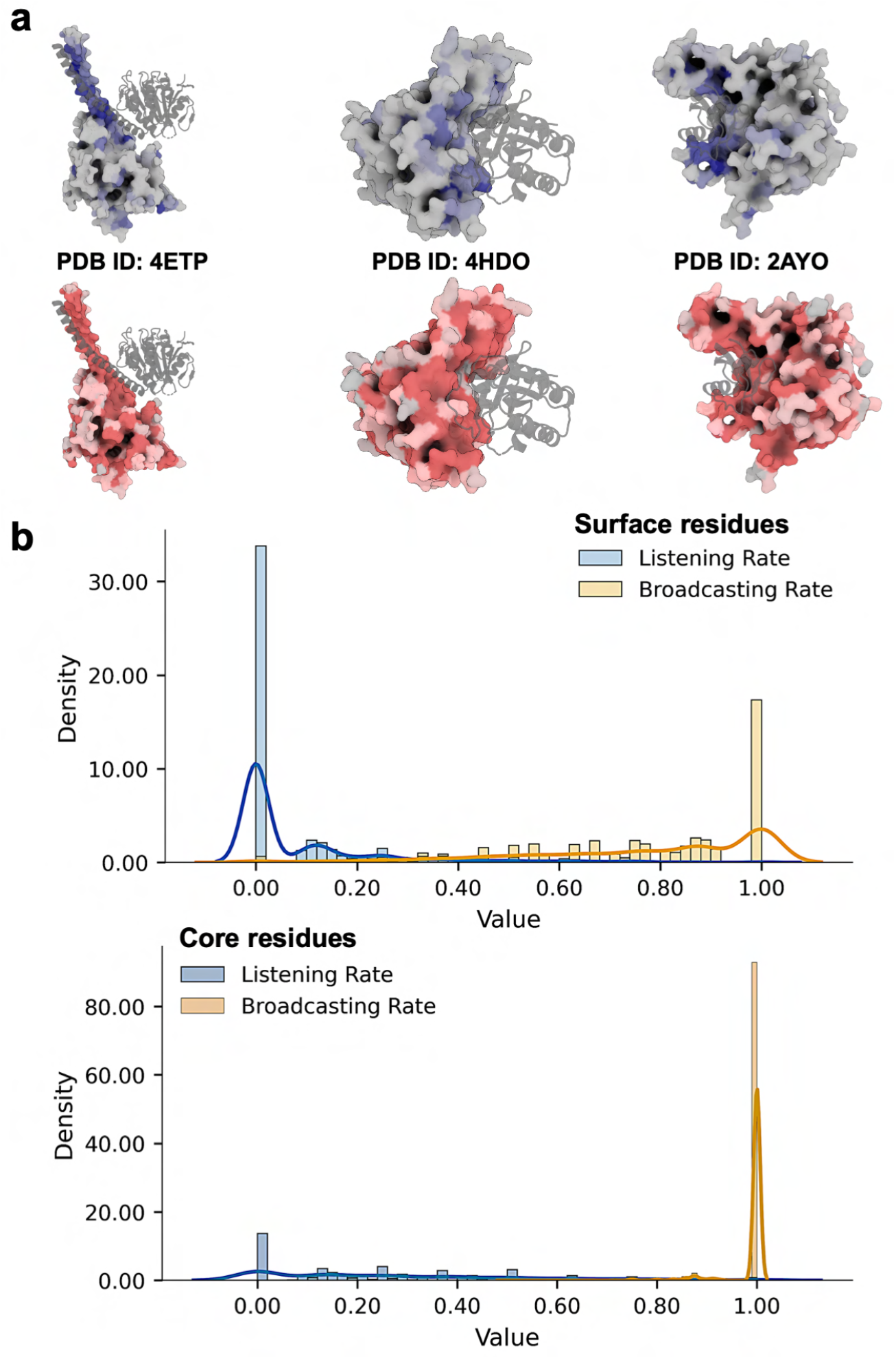
Listening and broadcasting for surface and core residues. **a)** visualized examples of listening (blue) and broadcasting (red) rate on protein surface **b)** The distribution of listening and broadcasting rate for two groups of core and surface residues.

**Figure S5:**
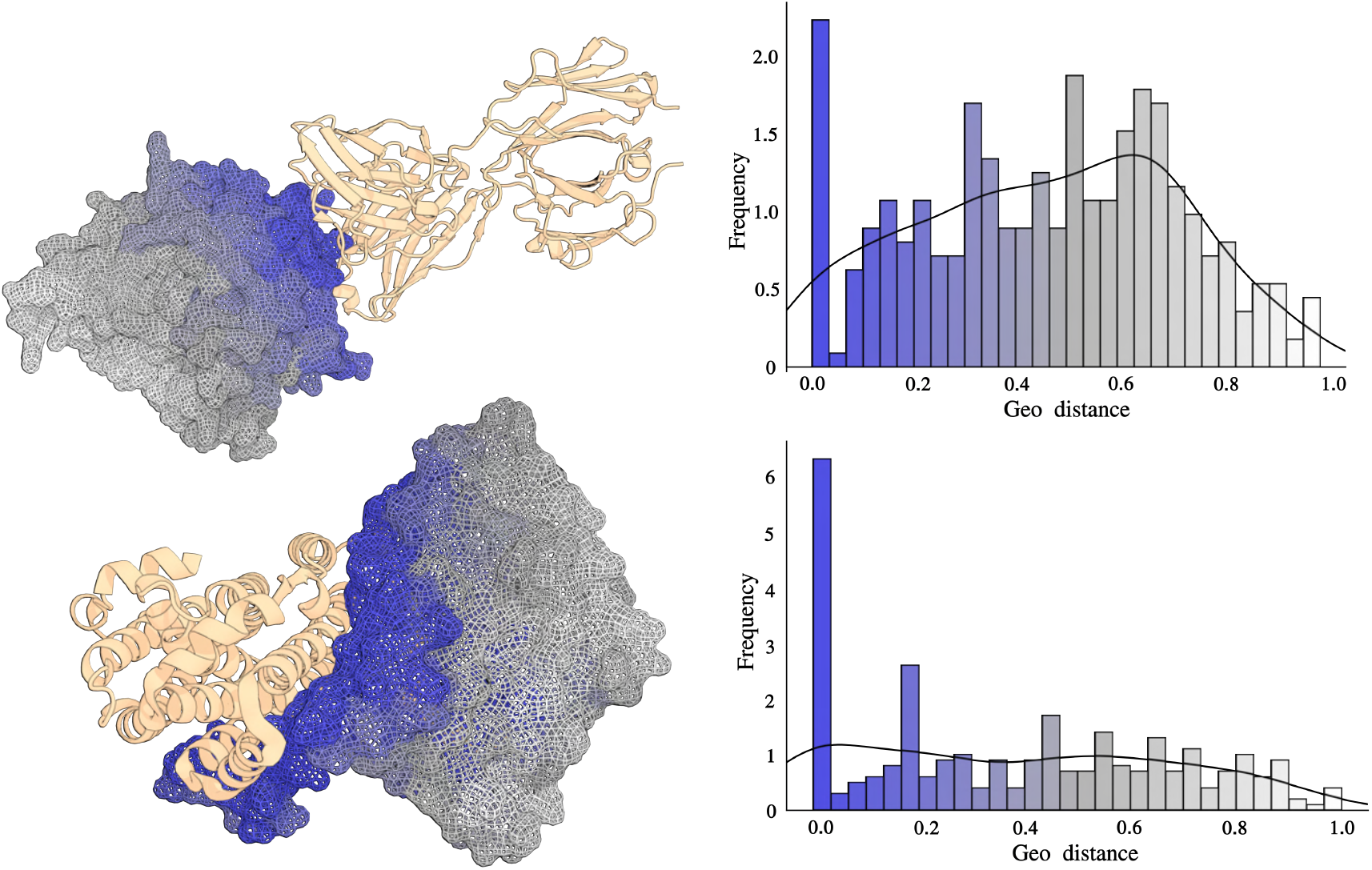
Geo Distance distribution. Two example of the distribution of residue level geo-distances with their corresponding visualization.

**Figure S6:**
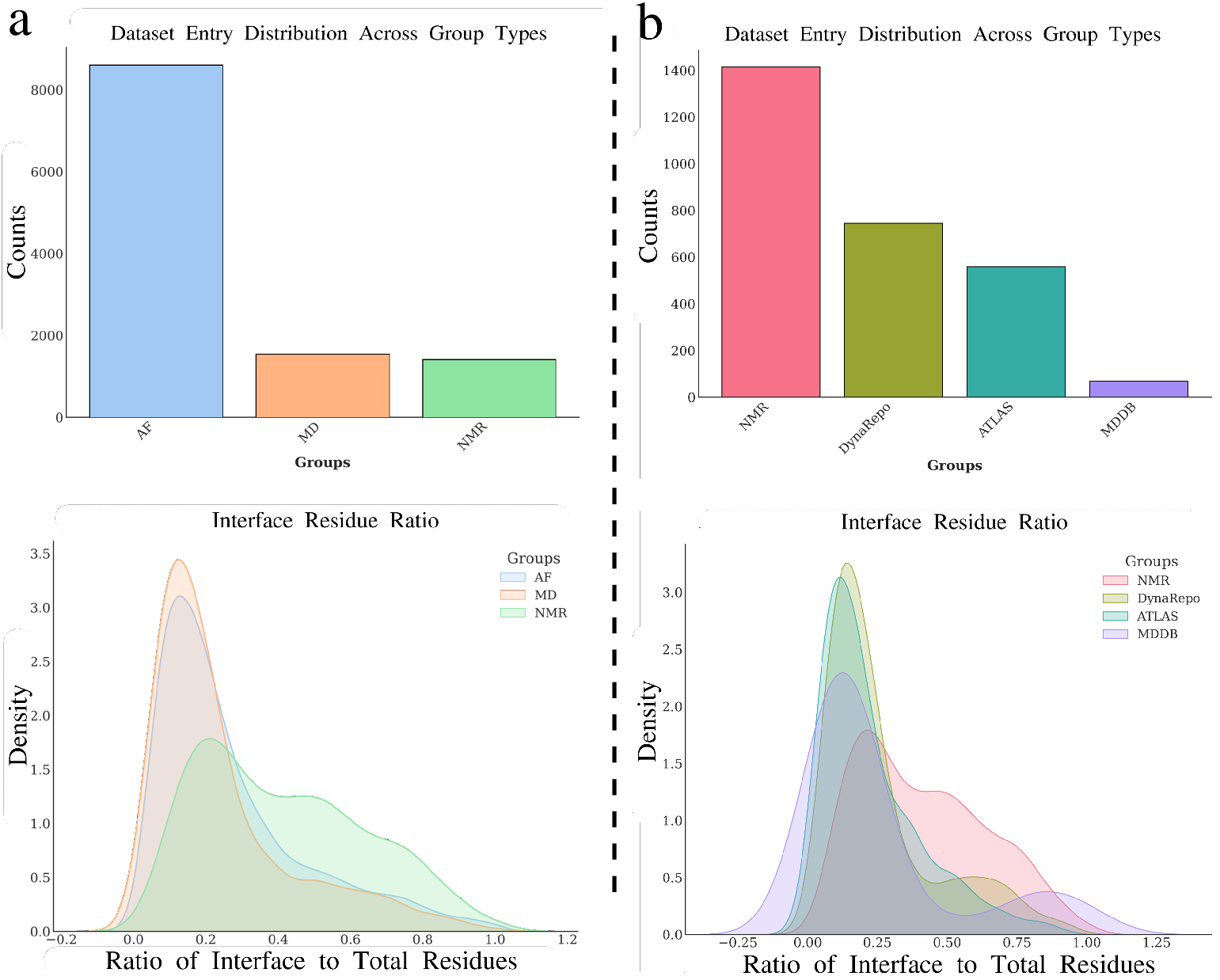
Dataset entry distribution across group types and the interface residue ratio analysis. **a)** The heterogeneity-aware model with all AlphaFlow (AF), MD simulation, and NMR entries. **b)** The dynamic-aware model with only NMR and MD simulation entries grouped by dataset source.

**Figure S7:**
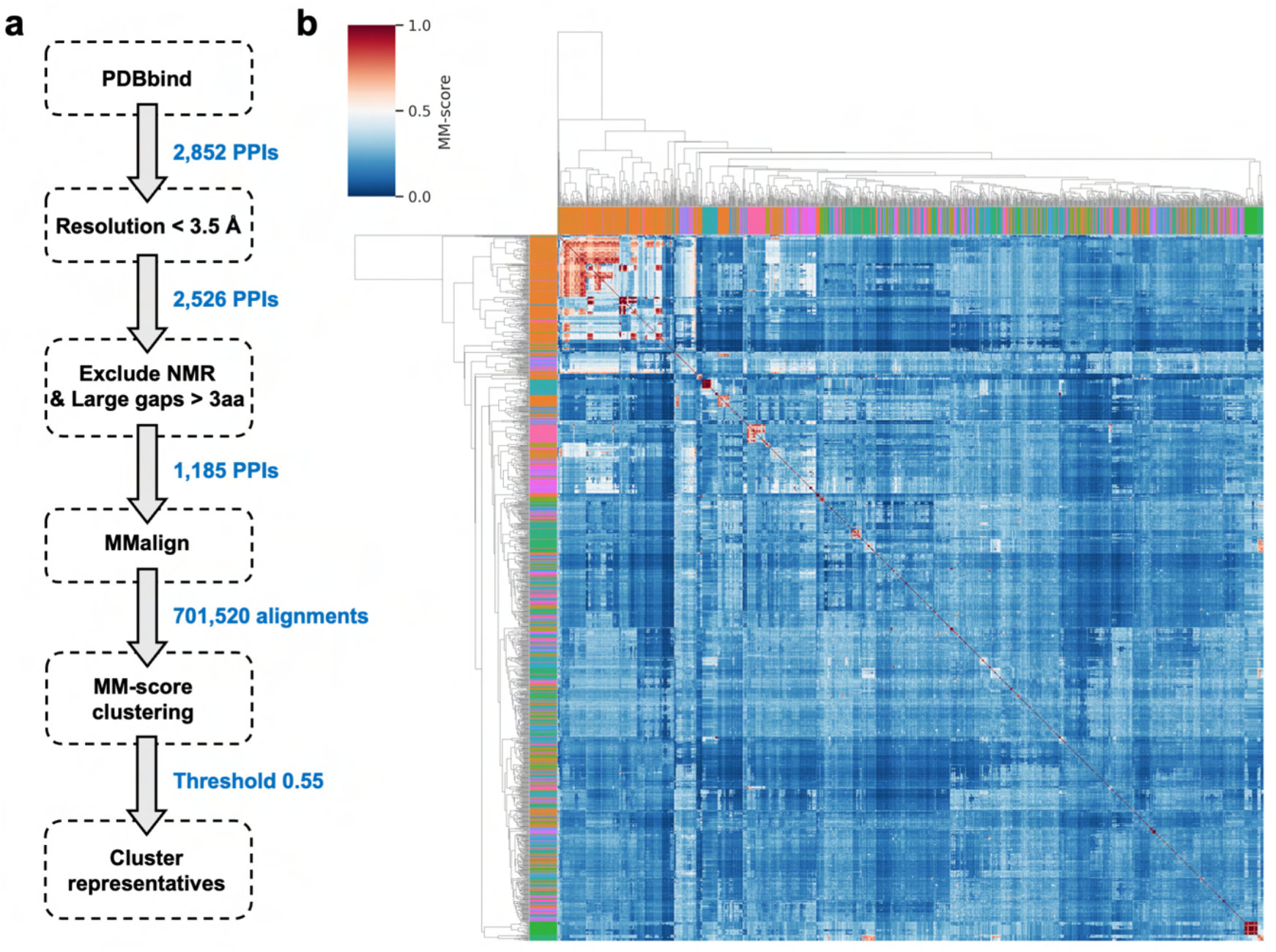
Building the MD simulation database from PDBbind. **a)** The general workflow shows how the PDBbind version 2020 is filtered based on resolution, and the number of gaps, and further clustered by performing pairwise alignment. **b)** Cluster heatmap displaying pairwise alignment results of representative structures, colored by MM-score.

**Figure S8:**
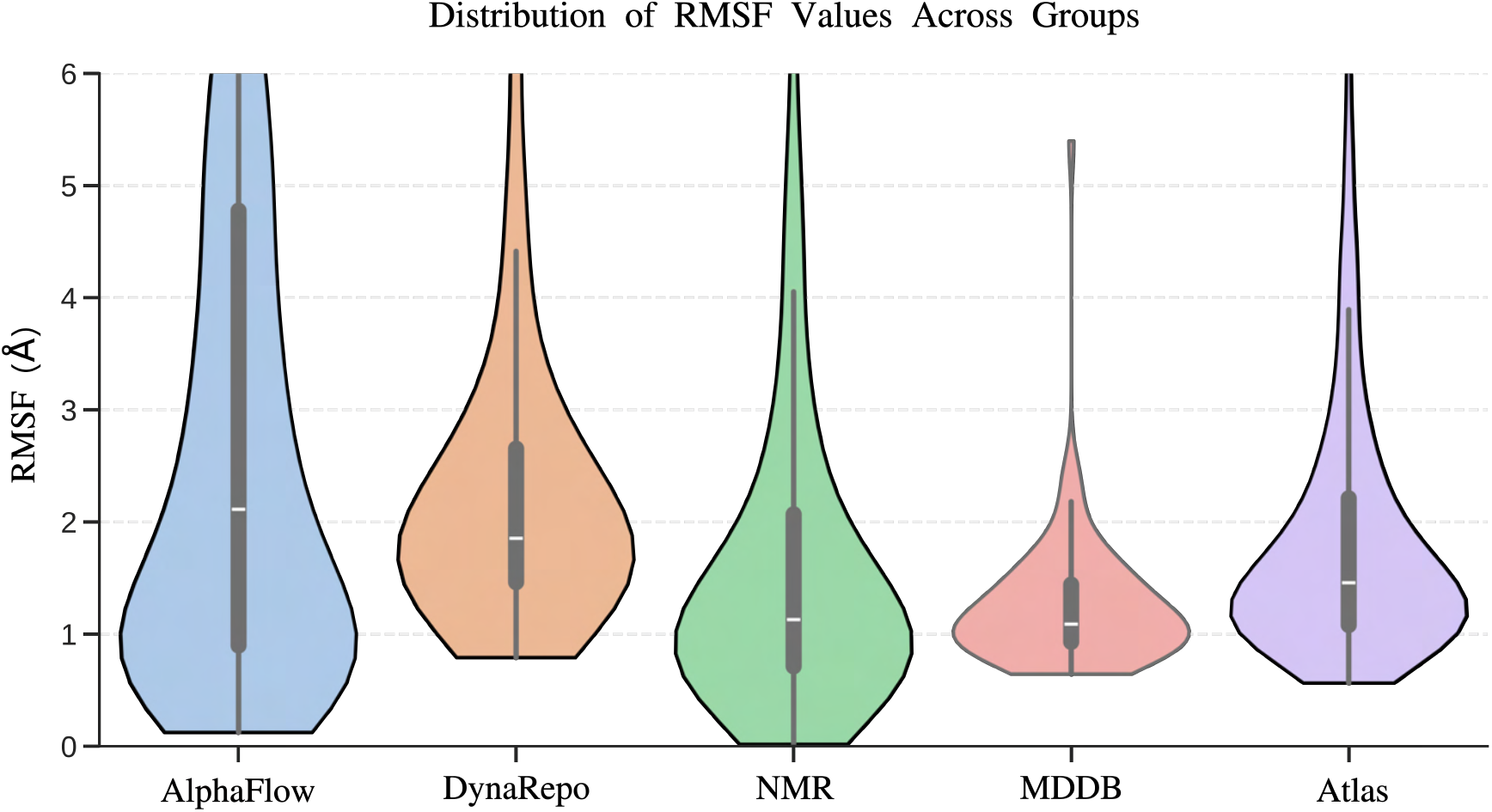
RMSF Across Data Source. Violin plot showing the RMSF distribution across dataset groups, excluding the outliers.

**Figure S9:**
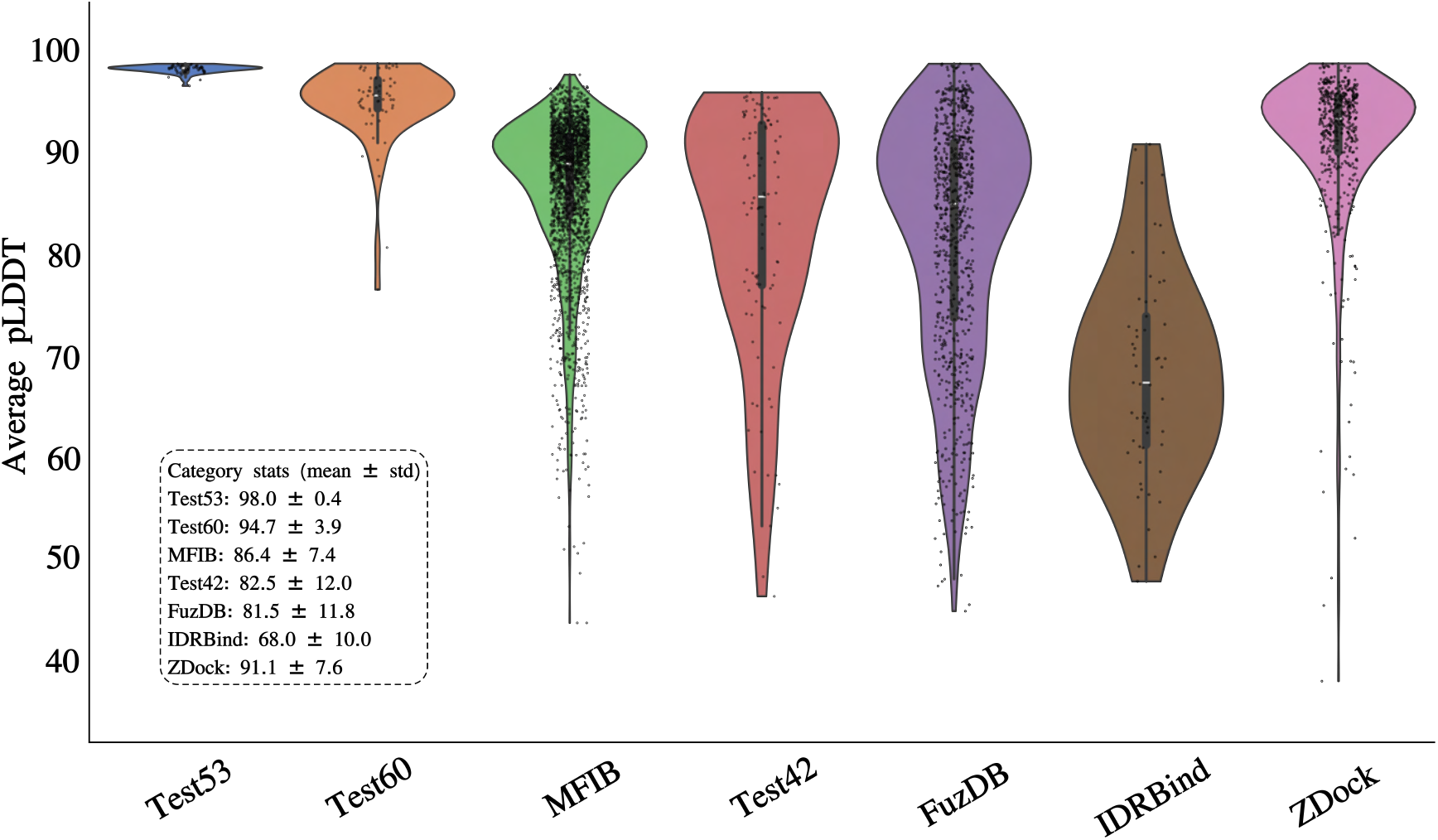
Violin plot of pLDDT distributions across different benchmark categories. The distribution of predicted Local Distance Difference Test (pLDDT) scores across various benchmark datasets. Individual data points are represented as scatter overlays, while the inner boxplots indicate the interquartile range and median.

**Figure S10:**
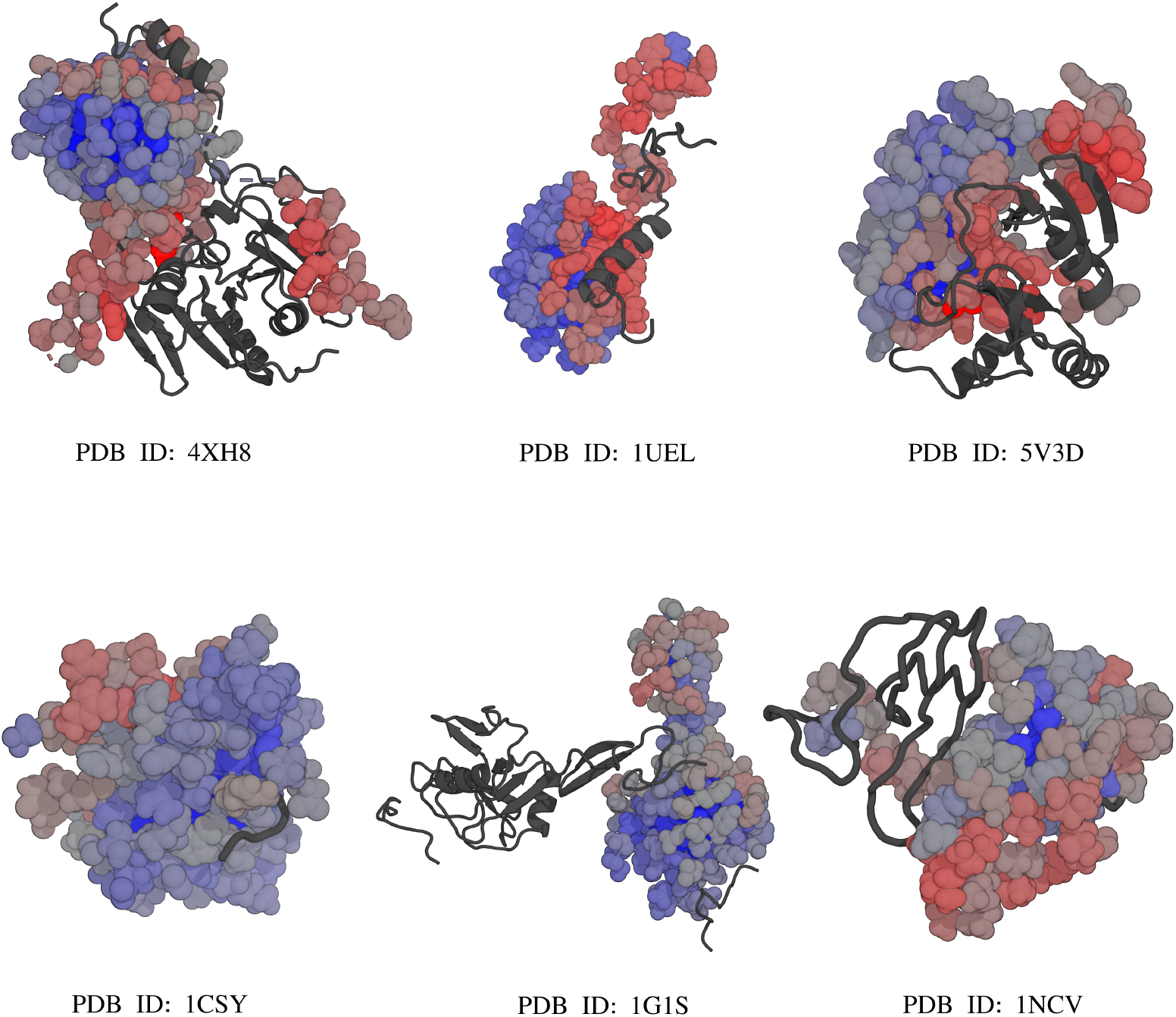
Examples of IDR structures. Selected examples illustrating good and poor predictions for intrinsically disordered regions.

**Figure S11:**
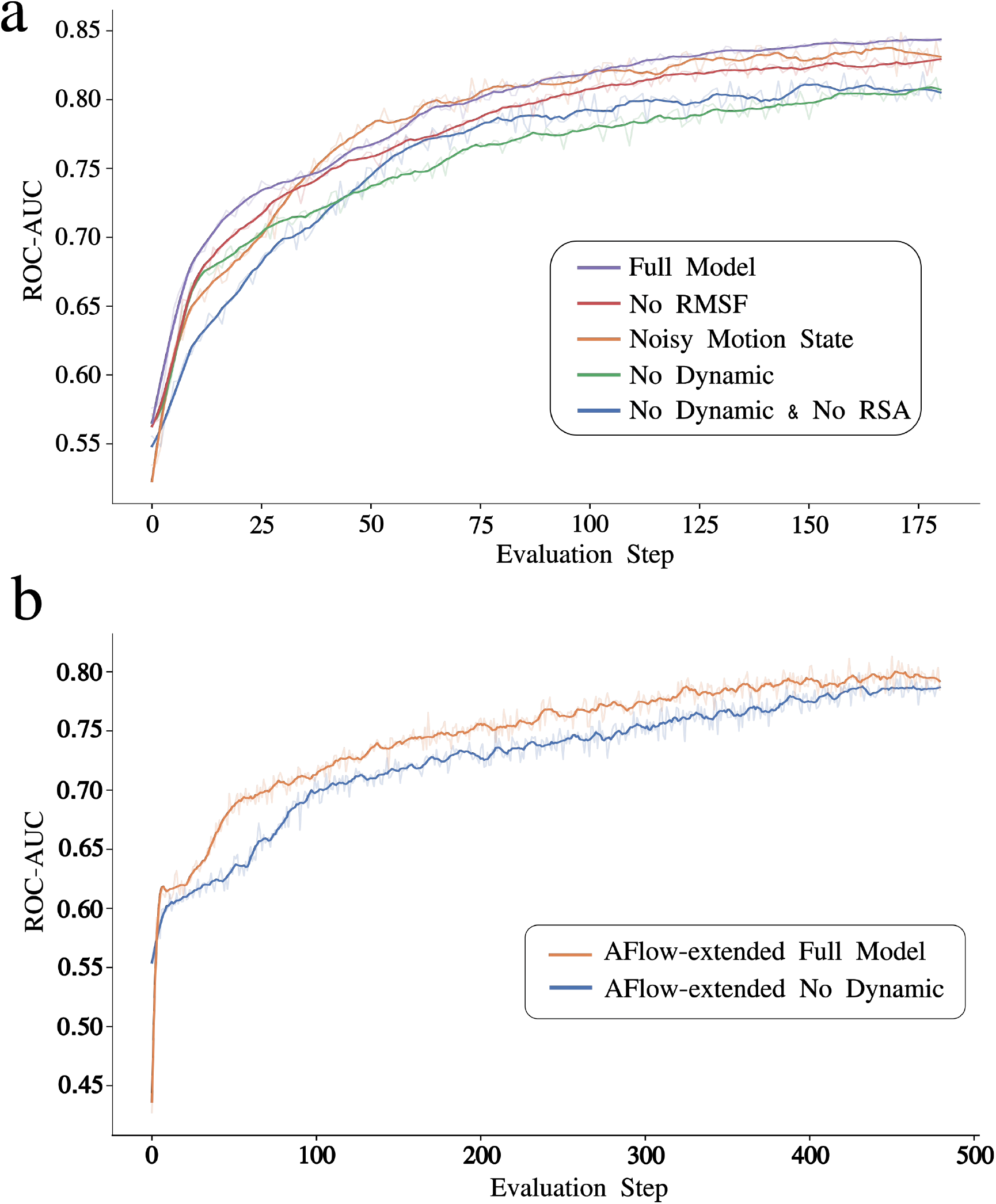
Ablation study on the impact of dynamic features and cooperative message passing. The line plot illustrates the validation set performance over time steps during the training process. **a)** represent the ablation study in details for different combination of dynamic features and cooperative message passing for dynamic-aware model trained over MD simulations and NMR data. **b)** Indicates the difference in ROC-AUC for the heterogeneity-aware model trained over all data including AlphaFlow conformations, with and without dynamic features.

**Figure S12:**
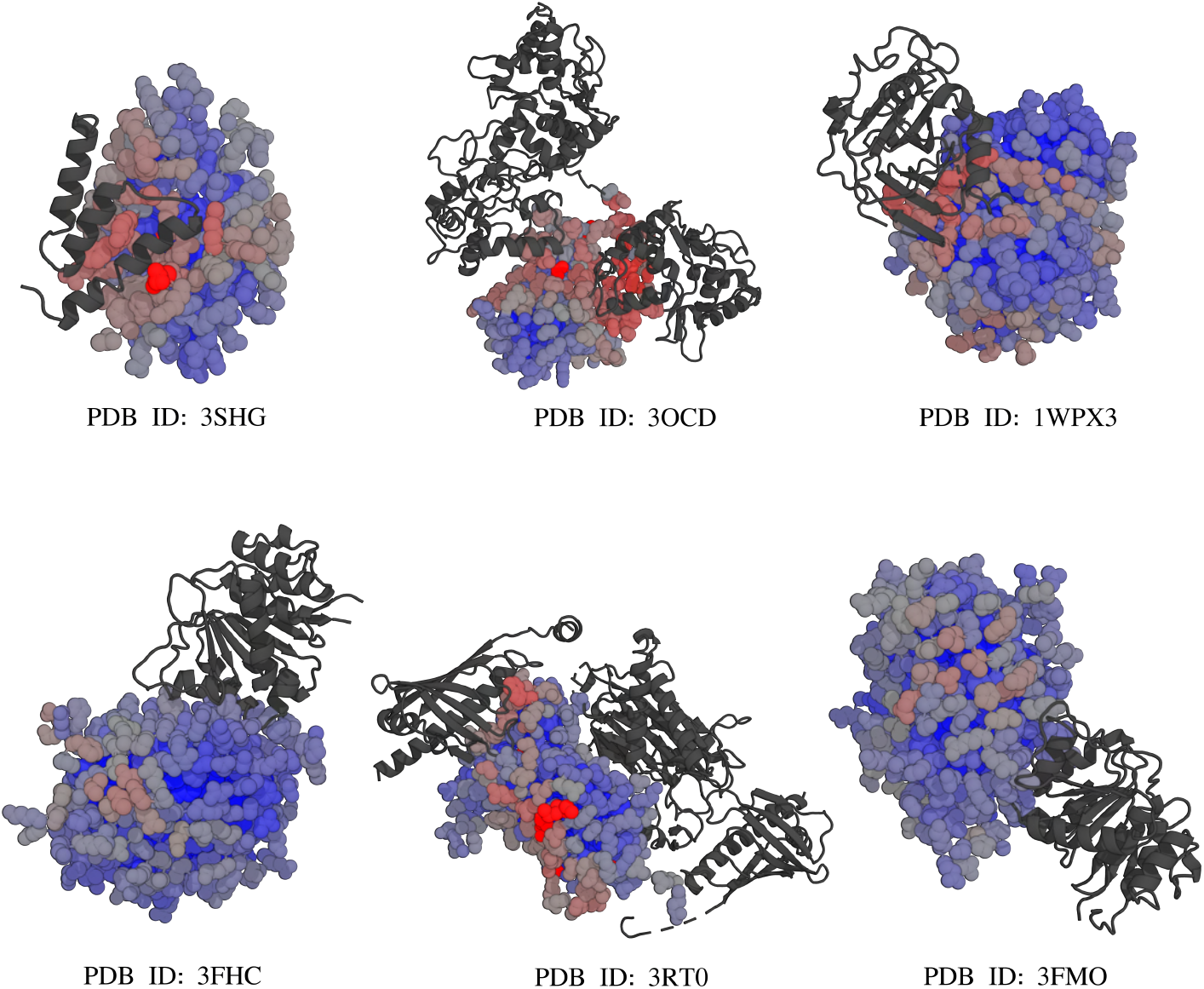
General structural examples. Selected cases showing good and poor prediction performance for static or transient complexes.

**Figure S13:**
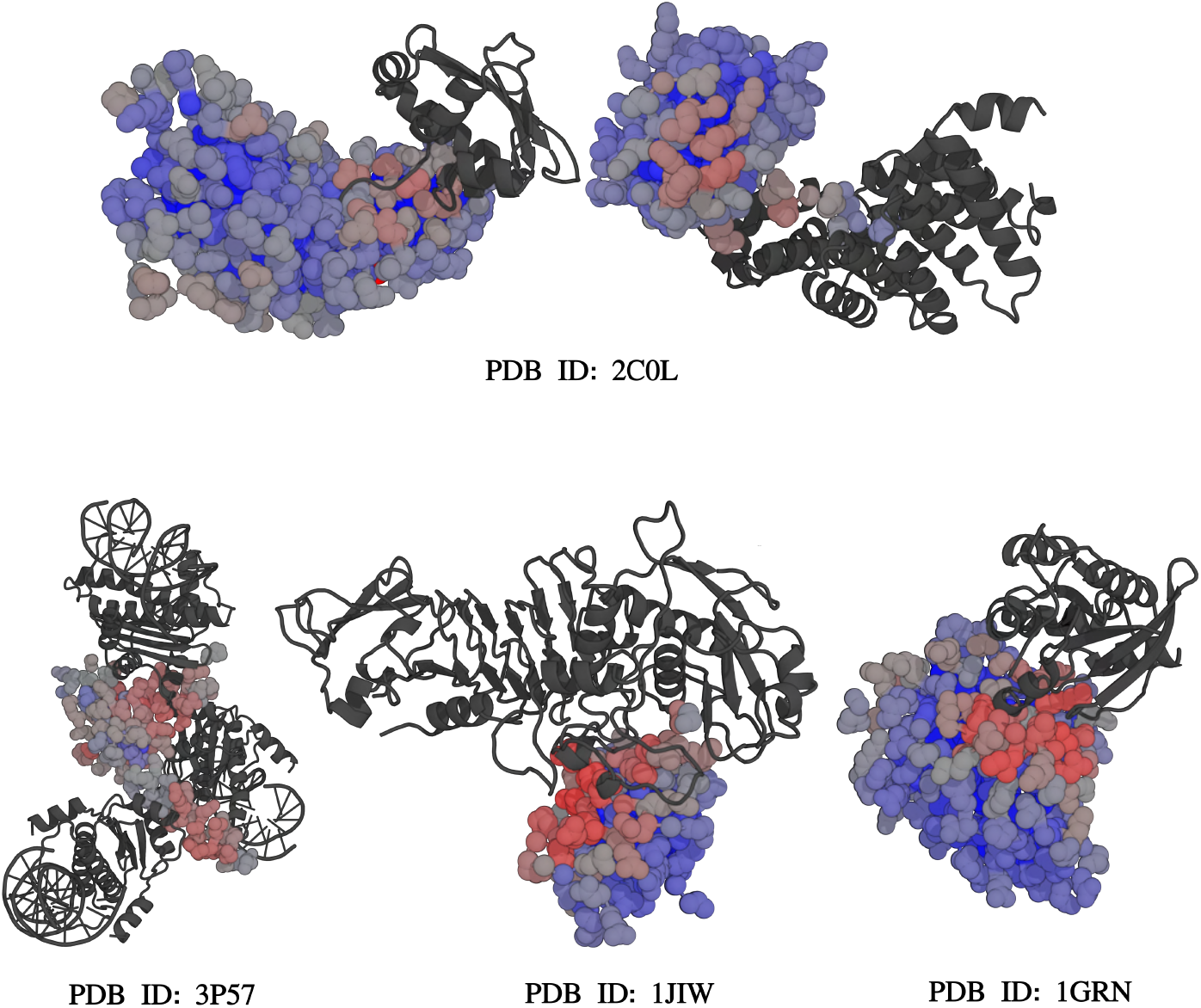
ZDock examples. Representative examples from ZDock benchmark.

## Supplementary Tables

**Table S1:** Number of unique chains per PDBbind entry before and after filtering steps (see Figure S7).

| Number of Unique Chains | Before Filtering | After Filtering |
| --- | --- | --- |
| 2 | 1608 | 347 |
| 3 | 637 | 49 |
| 4 | 165 | 6 |
| 1 or >4 | 112 | 7 |

**Table S2:** Median values of performance metrics for different datasets. Values are rounded to two decimal points.

| Dataset | Accuracy | Precision | Recall | Specificity | MCC | ROC-AUC | PR-AUC |
| --- | --- | --- | --- | --- | --- | --- | --- |
| MFIB | 0.76 | 0.64 | 0.92 | 0.67 | 0.52 | 0.88 | 0.83 |
| Test42 | 0.78 | 0.60 | 0.92 | 0.74 | 0.47 | 0.85 | 0.82 |
| ZDock | 0.80 | 0.41 | 0.50 | 0.86 | 0.28 | 0.79 | 0.46 |
| FuzDB | 0.67 | 0.56 | 1.00 | 0.12 | 0.26 | 0.77 | 0.77 |
| Testing transient | 0.87 | 0.38 | 0.42 | 0.92 | 0.32 | 0.85 | 0.41 |
| IDRBind | 0.71 | 0.65 | 1.00 | 0.20 | 0.44 | 0.79 | 0.84 |
| Test60 | 0.82 | 0.39 | 0.59 | 0.85 | 0.32 | 0.81 | 0.48 |

**Table S3:** Mean values of performance metrics for different datasets. Values are rounded to two decimal points.

| Dataset | Accuracy | Precision | Recall | Specificity | MCC | ROC-AUC | PR-AUC |
| --- | --- | --- | --- | --- | --- | --- | --- |
| MFIB | 0.74 | 0.63 | 0.87 | 0.62 | 0.50 | 0.86 | 0.81 |
| Test42 | 0.77 | 0.57 | 0.77 | 0.63 | 0.43 | 0.83 | 0.71 |
| ZDock | 0.79 | 0.46 | 0.54 | 0.82 | 0.35 | 0.81 | 0.53 |
| FuzDB | 0.65 | 0.54 | 0.90 | 0.27 | 0.27 | 0.75 | 0.71 |
| Testing transient | 0.86 | 0.36 | 0.42 | 0.91 | 0.31 | 0.82 | 0.42 |
| IDRBind | 0.68 | 0.66 | 0.97 | 0.28 | 0.41 | 0.75 | 0.80 |
| Test60 | 0.78 | 0.41 | 0.57 | 0.80 | 0.33 | 0.81 | 0.50 |

